# Siglec-7 and -15 recognize repeated clustered sulfo-sialo O-glycan motifs on select O-glycoproteins

**DOI:** 10.64898/2026.01.07.698156

**Authors:** Felix Goerdeler, Daniël L. A. H. Hornikx, Vera T. C. Valckx, Yu-Chun Chien, Dylan de Wit, Thapakorn Jaroentomeechai, Martin Jaeger, Sanae Furukawa, Huan-Chuan Tseng, Jaesoo Jung, Edward N. Schmidt, Kay-Hooi Khoo, Rebecca L. Miller, Wilfred T. V. Germeraad, Gerard M. J. Bos, Matthew S. Macauley, Henrik Clausen, Christian Büll, Yoshiki Narimatsu

## Abstract

Siglecs are key immunoreceptors in immune homeostasis and cancer immunosuppression. Diverse endogenous and exogenous sialoglycan ligands induce Siglec-mediated immunoregulation, but the nature of endogenous Siglec ligands and how seemingly ubiquitous sialoglycans achieve selectivity is unclear. Here, we employed a HEK293 cell-based array and glycoprotein reporters designed from human mucins and mucin-like proteins with O-glycans in natural clusters and repeat motifs to dissect specificities of Siglec-7 and −15. We first used precomplexed Siglec-Fc chimera to demonstrate that CHST1-mediated 6-*O*-sulfation markedly enhanced binding to select O-glycoproteins, namely MAdCAM1, CD43, and PSGL-1. Probing binding properties of Siglec-7 expressed exogenously on CHO cells or endogenously on human monocytes by reverse assays employing fluorophore-tagged O-glycoprotein reporters revealed specific binding to the sulfo-sialyl core1 O-glycoform of the same O-glycoproteins. Our study indicates that Siglec-7 on immune cells employs multivalent interactions with repeated motifs of clustered sulfo-sialyl O-glycans to drive selectivity in ligand interactions and Siglec-induced immunosuppression.

## INTRODUCTION

Immunoregulatory glycan-binding proteins (GBPs) are key receptors for self-recognition and maintaining immune homeostasis. One important class of immunoregulatory GBPs are Siglecs (sialic acid-binding immunoglobulin-like lectins) which recognize sialic acid residues on glycans expressed by host cells and mainly induce immunosuppressive activities ^1^. There are 14 functionally expressed human Siglecs and these share a common structure with the extracellular domain comprising an N-terminal V-set immunoglobulin (Ig)-like domain that mediates sialoglycan binding followed by varying numbers of C2-set Ig-like domains that position the V-set domain away from the cell surface ^2^. The V-set Ig-like domain contains a conserved arginine residue that binds to sialic acid via an essential salt bridge and a highly variable CC’ loop determining ligand specificity ^3–5^. The intracellular domain contains an immunoreceptor tyrosine-based inhibitory motif (ITIM), or in the case of Siglec-14/15/16 charged residues that recruit adaptor proteins (DAP10/12) carrying immunoreceptor tyrosine-based activation motifs (ITAMs) ^2,6,7^. Upon binding to specific sialoglycans expressed by host cells, Siglecs cue immunomodulatory signals via their ITIM/ITAMs. For instance, Siglec-2 (CD22) serves in inhibition of B-cell activation, whereas Siglec-7 and Siglec-15 serve in suppression of NK cell and macrophage responses, respectively ^8–10^. The immunomodulatory effects of Siglecs are exploited by cancer cells and invading pathogens by expression of related sialoglycans to engage Siglecs and evade immune responses ^2,11^. The use of aberrant sialoglycans by cancer cells to suppress immunity renders Siglecs promising immune checkpoints for therapeutic intervention ^12–15^. Despite the key roles of Siglecs in immune regulation and diseases, our understanding of how natural sialoglycan ligands confer sufficient specificity to support Siglec function is still limited ^2,16^.

Part of the challenges with identifying ligands for Siglecs lies in experimental limitations. Most studies of the binding specificities of Siglecs have so far mainly relied on binding to oligosaccharides and/or neoglycolipids, where screening of large libraries of oligosaccharides by printed glycan arrays and neoglycolipid arrays has provided valuable insights into the glycan binding properties of different Siglecs ^17–24^. However, individual oligosaccharides do not necessarily provide the natural context in which glycans are found on proteins and lipids at the cell surface. Contextual presentation of sialoglycans on proteins has been proposed as a mechanism to increase diversity of ligands, yielding sufficiently unique, high-avidity ligands to orchestrate regulation of Siglec function ^1,25–29^. Glycoproteins may contain multiple glycans in repeat motifs and/or clusters that are arranged in ways to support multivalent interactions with the predicted oligomeric states in which Siglecs are found on cells. For example, mucins and some mucin-like glycoproteins characteristically contain repeated O-glycan motifs in extended O-glycodomains ^30^. Probing for such contextual interactions with glycans on proteins is possible with the cell-based glycan array platform that can display the natural context of glycans on glycoproteins and glycolipids ^31,32^. Cell-based arrays consist of libraries of isogenic cells with engineered glycosylation capacities through combinatorial KO/KI of glycosyltransferase genes, where individual cells display distinct glycan features in context of proteins and lipids on the cell surface for interrogation and dissection of glycan-binding interactions ^33^. We previously used cell-based glycan arrays to demonstrate that Siglec-7 and −15 bind to di-sialylated core1 (dST) and sialyl-Tn (STn) O-glycans, respectively ^34^, as precomplexed oligomeric Fc fusion proteins, in line with other studies ^10,35,36^. Moreover, studies show that Siglec-7/15 binding is markedly enhanced by CHST1-mediated 6-*O*-sulfation of galactose ^34,37–39^. The sulfated ligands of Siglec-7 are mainly O-glycans whereas Siglec-15 can recognize both O- and N-glycans when sulfated ^34,37,38^. In addition, we found evidence that the O-glycan carrier proteins influence Siglec binding ^34^. For example, despite its preference for dST ligands, Siglec-7 could also bind to STn O-glycans when presented in the context of mucins ^34^, and several studies have identified particular glycoproteins as targets for Siglec binding. A genome-wide CRISPR screen in K562 leukemia cells identified the mucin-like protein CD43 as a main O-glycoprotein ligand for Siglec-7 ^40^. Subsequent biochemical characterization revealed that a stretch of dense O-glycan clusters presented on the N-terminal region of CD43 appeared to be important for Siglec-7 binding ^40,41^. In another study using JVM-3 leukemia cells, not only CD43, but also CD45 and PSGL-1 were identified as Siglec-7 ligands ^35^, whereas a recent study in DLD-1 colon cancer cells identified the two O-glycoproteins MUC13 and PODXL as Siglec-7 ligands ^42^. Specific glycoprotein ligands have also been determined by immunoprecipitation for other Siglecs, e.g. Siglec-3 and −8 ^43–45^, clearly supporting the hypothesis that the protein context, i.e. the arrangement of glycans on proteins in clusters or patterns, has a profound impact on Siglec recognition.

More than 30 years ago, Ajit Varki proposed that simple sialoglycan epitopes could not support binding and orchestrate the diverse biological functions of Selectins and Siglecs simply because the structural variation of mammalian glycan binding epitopes is too limited ^25^. Instead, more complex and selective interactions were envisioned through “clustered saccharide patches” (CSPs) comprised of multiple discontinuous glycan moieties as well as multivalency through oligomeric states of GBPs as mechanisms to enhance specificity ^1,25–28^ (**Fig. 1**). The structural molecular nature of CSPs and how such may be recognized is still unclear, but the typical clustering of O-glycans on mucins and O-glycoproteins was considered one likely scenario ^25,26,46^ (**Fig. 1a**). The extent to which the binding site(s) of individual receptor proteins and/or their oligomeric assemblies and arrangement in cell membranes direct binding is also poorly understood ^25,26^. In this respect, it is relevant to note that the detection of Siglec binding to glycans has often required precomplexing of Siglec-Fc chimera with an IgG antibody or StrepTactin ^21,47^, and this may bias for multivalent interactions and potentially binding to unnatural arrangements (clustering) of glycans (**Fig. 1b**). At the cell surface, the organization of Siglecs remains to be elucidated but may involve dynamic assembly of oligomers by exposure to ligands ^29,50,51^. While Siglecs can interact with adjacent sialoglycans at low affinity, a process termed *cis*-masking, the oligomeric state and dynamics may change upon exposure to high-affinity and/or high-avidity *trans*-ligands ^2,52,53^ (**Fig. 1c**). The O-glycan-binding Siglec-7 and −15 are obvious candidates to explore and dissect with these parameters, and currently these receptors are believed to have receptor binding sites that only accommodate a simple sialic acid-capped glycan moieties on O-glycoproteins ^54,55^. However, Siglec-7 is reported to bind the disialogangliosides GT1b, GD2, and GD3 ^3,5,23,56–59^, and more recently, the presence of an additional sialic acid binding site on Siglec-7 has also been proposed to accommodate binding to two sialic acid residues in gangliosides ^60^.

**Figure 1.**
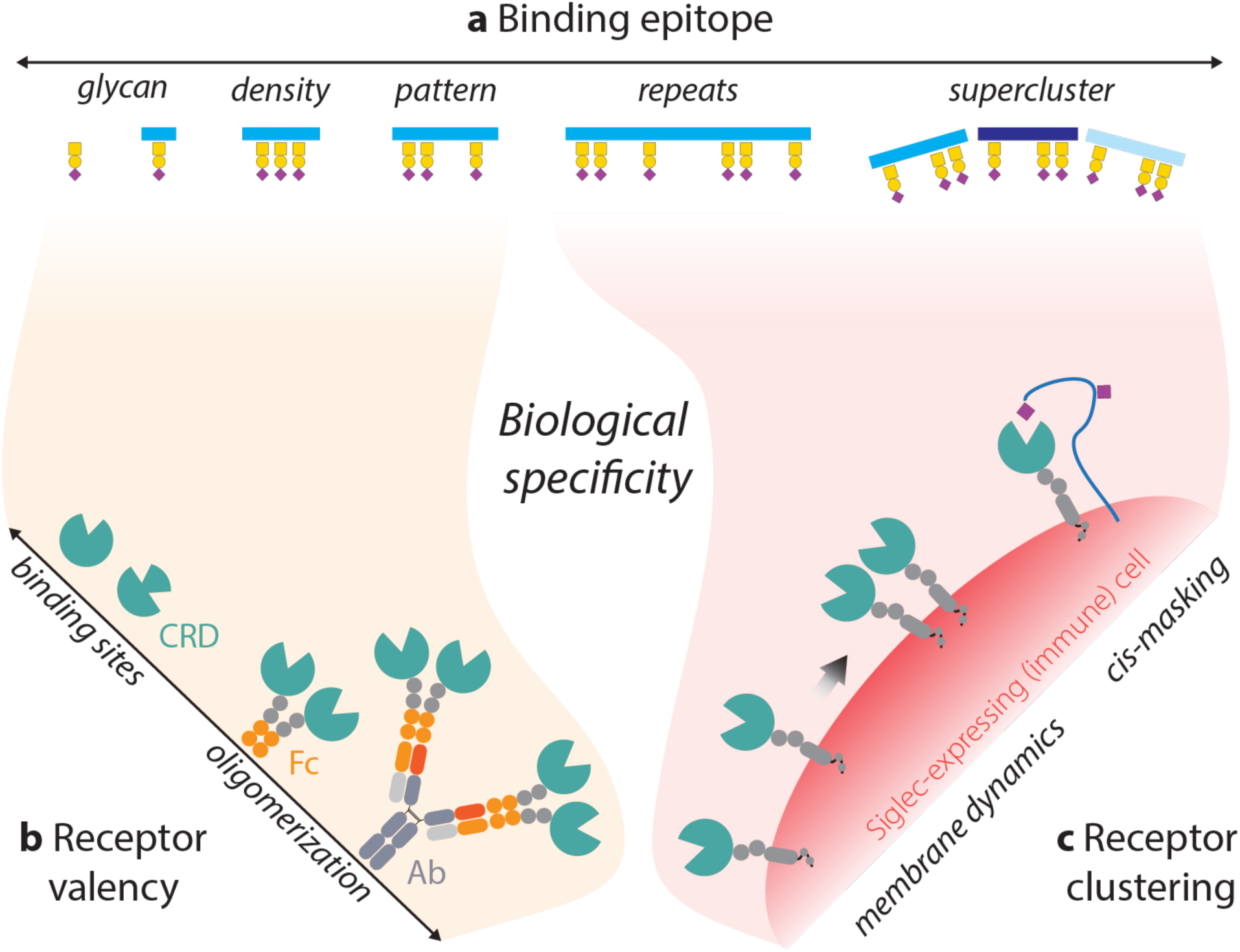
Clustered saccharide patches (CSPs) formed by contextual O-glycan arrangements, as well as oligomerization of binding receptors potentially drive specificity, illustrated with Siglecs. CSPs, also considered discontinuous glycan epitopes, are conceptionally defined as binding motifs for glycan-binding proteins (GBPs) that are formed by multiple glycans and go beyond simple density effects of high glycan concentrations ^1,25–27^. CSPs are proposed to expand the sialoglycan epitome and provide for more selective and high-avidity interactions. However, to dissect high-avidity binding to CSPs, the valency and spacing/orientation of glycan ligands leading to potential recognition patterns as well as the surface distribution and oligomeric modes of receptors need to be scrutinized. (**a**) **Binding epitope** – The characteristic clustering of O-glycans in distinct patterns (often found in tandem repeat regions of mucins and mucin-like domains of O-glycoproteins) is perhaps the most obvious model for CSPs, with one simple example being clusters of 1-3 adjacent O-glycans. Indeed, antibodies to truncated Tn and STn O-glycans often selectively bind two adjacent O-glycans ^95–97^. However, more contextual patterns and repeats of multiple O-glycan clusters, either on the same protein (such as shown in this study) or on different proteins (“superclusters”), may also be considered. (**b**) **Receptor valency** – The carbohydrate-recognition domain of Siglecs may have one or more sialic acid binding sites ^60,67^ and direct binding studies with isolated recombinant Siglecs often require artificial oligomeric assemblies for detection of binding, e.g., Fc chimera and precomplexing of these, which may affect multivalent binding (shaded in orange). (**c**) **Receptor clustering –** The natural oligomeric state of Siglecs at the cell surface and potential dynamic changes in response to ligand exposure are poorly understood. Moreover, the ligand binding sites of Siglecs in the cell membrane may be preoccupied by interactions with adjacent sialic acid ligands, referred to as *cis*-masking ^52,53^. Binding to Siglecs may thus only be detectable upon exposure to ligands with higher affinity and/or avidity, which may involve changes in oligomeric states. In conclusion, higher-order presentation of the glycan ligands as well as of the receptors (shaded in red) are key variables to consider in the natural context, likely conveying biological specificity to Siglecs. Figure was inspired by original concepts proposed by Ajit Varki ^1,25–27^.

Here, we show that contextual presentation of sulfo-sialo O-glycans does indeed guide binding of Siglec-7 and - 15 to select mucin-like O-glycoproteins. We employed libraries of glycoengineered cells to display and produce O-glycoproteins with custom-designed O-glycans and dissect contextual interactions by Siglecs. We used cell-based mucin arrays with engineered O-glycan sialylation and sulfation to demonstrate that precomplexed Siglec-7 preferentially binds 6-*O*-sulfated O-glycans with α2-6Sia (sulfo-dST/mSTb), while Siglec-15 binds 6-*O*-sulfated core1 with α2-3Sia independent of α2-6Sia (sulfo-mSTa/dST). Employing an expanded library of reporters designed from mucin-like O-glycoproteins and displayed on glycoengineered cells, we identified the mucin-like O-glycoproteins MAdCAM1, CD43, and PSGL-1 as select sulfo-sialo O-glycoprotein targets for Siglec-7 and −15. We then isolated the corresponding secreted O-glycoprotein reporters and used these fluorophore-tagged in reverse assays to demonstrate that only the sulfo-dST glycoform supports binding to Siglec-7 exogenously expressed on CHO cells. We identified common sequence features among the MAdCAM1, CD43, and PSGL-1 O-glycoproteins that appear to direct select CHST1-sulfation of O-glycan clusters and support Siglec-7 binding to repeated saccharide patches in clustered O-glycans by multivalent interactions. Finally, to our knowledge, we demonstrate for the first time specific binding of these natural sulfo-sialo O-glycoprotein mimics to human monocytes that endogenously express Siglec-7. Together, our results support the concept that Siglec-7 employs contextual interactions with repeated clustered motifs of O-glycans in specific proteins to orchestrate selectivity in ligand recognition of sialoglycans.

## RESULTS

### Sulfated mucin-like O-glycoproteins are potent ligands for Siglec-7 and Siglec-15

The binding properties of human Siglec-7 and Siglec-15 (recombinant precomplexed Fc-chimera, hereafter Siglec-7/15-Fc) previously determined by our cell-based glycan and mucin arrays are depicted in **Figure 2a** ^31,34^, highlighting that (i) Siglec-7 can interact with STn when presented on mucins, (ii) Siglec-15 can interact with dST in context of CHST1-mediated 6-*O*-sulfation, and (iii) binding of both Siglec-7 and −15 is substantially boosted by 6-*O*-sulfation. To further dissect the basis for these interactions, we expanded on our genetic engineering of the glycosylation capacities in HEK293 cells to display additional structural variations of O-glycans. Using this expanded panel of glycoengineered HEK293 cells that displays O-glycans on endogenous membrane proteins, we found that Siglec-7-Fc binding was slightly enhanced by the loss of core2 O-glycans (HEK^KO:GCNT1^) and strongly boosted by introduction of CHST1 6-*O*-sulfation (HEK^KO:GCNT1;KI: CHST1^) (**Fig. 2b, Supplementary Fig. 1**). Interestingly, loss of α2-6-sialylation on dST O-glycans (HEK^KO:GCNT1/ST6GALNAC2–4^), which results in mSTa O-glycans, fully abrogated binding, similar to the total loss of sialylated O-glycans (HEK^KO:C1GALT1^). However, in the presence of CHST1 sulfation, the loss of α2-6-sialylation (HEK^KO:GCNT1/ST6GALNAC2-4;KI:CHST1^) markedly reduced but did not fully abrogate binding.

**Figure 2.**
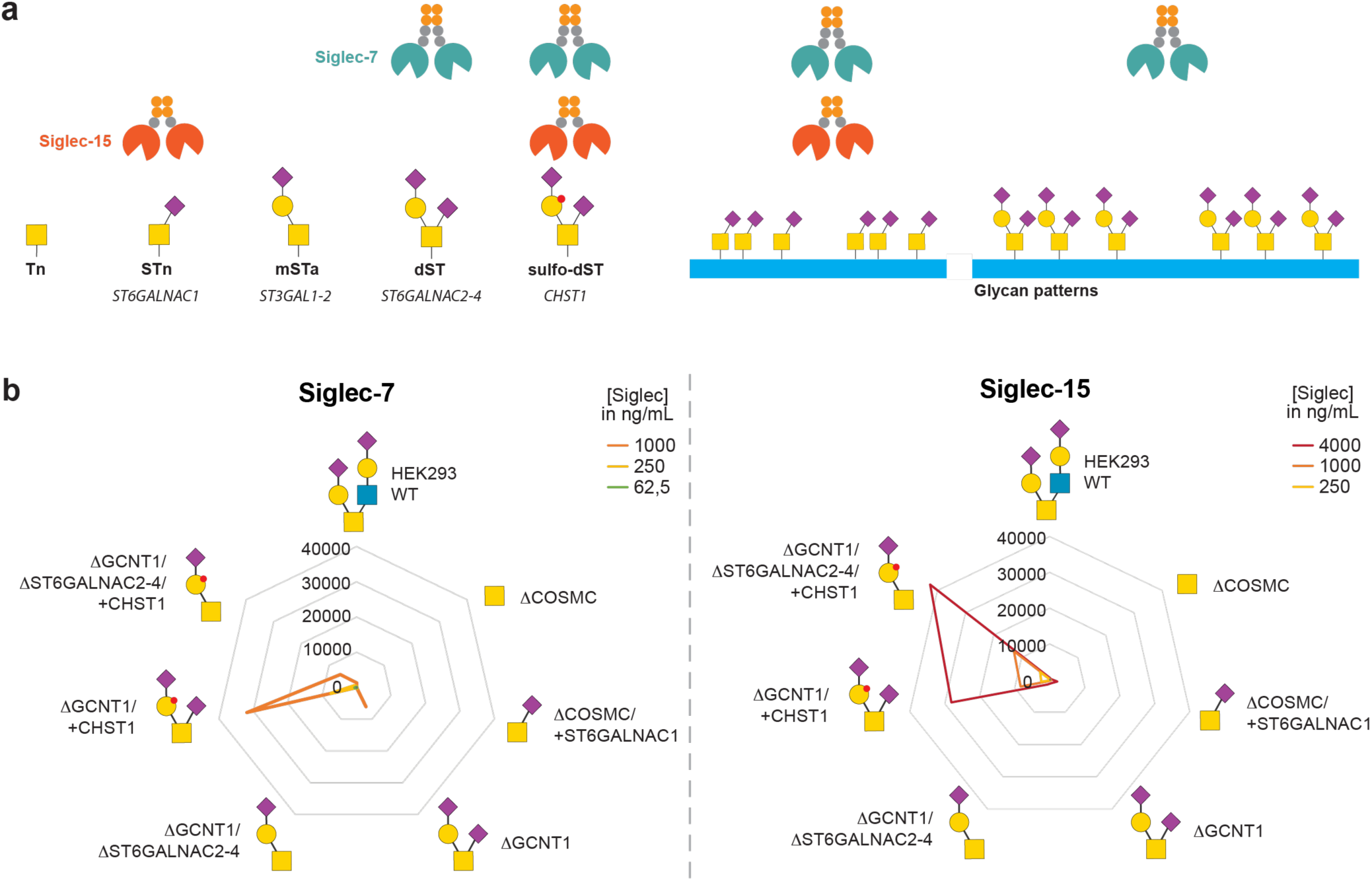
Genetic dissection of Siglec-7/15 binding to sulfated O-glycans on HEK293 cells. (**a**) Graphic depiction of O-glycan binding specificities of Siglec-7 and 15, showing that Siglec-7 can bind STn but only in context of clusters found on mucins. Glycan structures indicate common core1/2 O-glycans and follow the Symbol Nomenclature for Glycans (SNFG) ^98^. (**b**) Radar charts show flow cytometry analysis (mean fluorescence intensity, MFI) of Siglec-7/15 binding to glycoengineered HEK293 cells with KO/KI of glycosyltransferases and CHST1 sulfotransferase as indicated. Siglec-7/15 were precomplexed with anti-human-IgG antibody-Alexa647 (Invitrogen A-21445) and titrated to cells at the indicated concentrations. Representative MFI values from five independent experiments. Same data with CHST1 KI cell lines excluded is displayed in **Supplementary Fig. 1** to improve the resolution in the low MFI range.

Testing the same panel of cells with Siglec-15-Fc showed weak binding to STn (HEK^KO:C1GALT1;KI:ST6GALNAC1^) and dST O-glycans (HEK^KO:GCNT1^) and no binding to HEK^WT^ cells (**Fig. 2b, Supplementary Fig. 1**). Upon introduction of CHST1-mediated 6-*O-*sulfation (HEK^KO:GCNT1;KI:CHST1^), Siglec-15-Fc binding was increased 28-fold, and in contrast to Siglec-7-Fc, the binding was not reduced by loss of α2-6-sialylation of the inner αGalNAc residue (HEK^KO:GCNT1/ST6GALNAC2-4;KI:CHST1^) (**Fig. 2b**). Thus, both Siglec-7 and −15 bound efficiently to sialylated and 6-*O*-sulfated core1 O-glycans. However, while Siglec-7 binding was relatively dependent on the inner α2-6Sia residue, binding of Siglec-15 was completely independent.

Next, we elaborated on our previous finding that the context of O-glycans in clusters and patterns on mucins markedly influenced the binding of Siglec-7/15 ^34^. We transiently expressed a panel of reporter proteins (fused to GFP) containing representative sequences from human mucin tandem repeats (TRs) and mucin-like O-glycodomains of O-glycoproteins (collectively referred to as mucin reporters), on the cell surface or as secreted glycoproteins in glycoengineered HEK293 cells (**Fig. 3a, Supplementary Table 1**). To determine the binding increase upon expression of the mucin reporters, we subtracted MFIs of parent glycoengineered cells from MFIs of mucin-expressing cells using the GFP fluorescence of the mucin reporters to distinguish cell populations. Furthermore, we validated select interactions with secreted, purified mucin reporters by ELISA.

**Figure 3.**
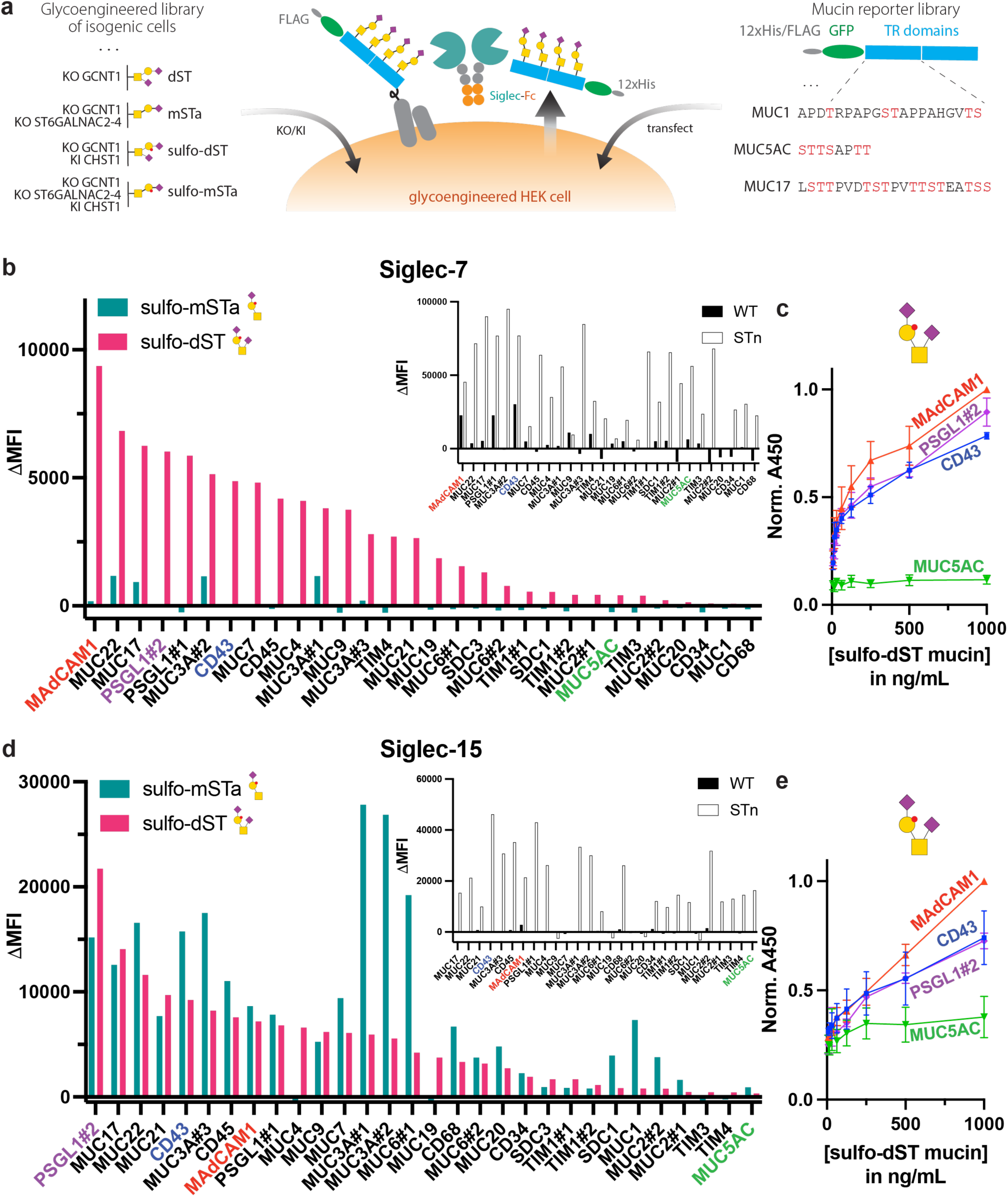
Siglec-7/15 binding to sulfated O-glycans on HEK293 cells expressing mucin reporters. (**a**) Depiction of the cell-based platform for display and production of mucin reporters with defined O-glycans. The glycoengineering designs for sulfated ST O-glycans are indicated, and the general reporter design with GFP, tags, and mucin O-glycodomains is illustrated. Sequences of the mucin glycodomains included in the reporter library are shown in **Supplementary Table 1.** (**b**) Flow cytometry analysis of Siglec-7 (250 ng/mL) binding to cells engineered with CHST1 sulfation and dST or mSTa O-glycosylation capacities, expressing mucin reporters as indicated. Mucin reporters were ranked by their relative increase in Siglec-7 binding upon mucin expression (ΔMFI), and mucin reporter expression was monitored by GFP signal. Inset: Comparative analysis of our previous data from ^34^, showing Siglec-7 binding increase (ΔMFI) upon mucin expression in HEK^WT^ or STn cells (HEK^KO:COSMC;KI:ST6GALNAC1^). **(c)** ELISA analysis of Siglec-7 binding to select purified recombinant mucin reporters with sulfo-dST O-glycans; values represent mean ± SEM from n = 3. **(d)** Flow cytometry analysis of Siglec-15 (1 µg/mL) binding to cells engineered with CHST1 sulfation and dST or mSTa O-glycosylation capacities, expressing mucin reporters as indicated. Mucin reporters were ranked by their relative increase in Siglec-15 binding upon mucin expression (ΔMFI), and mucin reporter expression was monitored by GFP signal. Inset: Comparative analysis of our previous data from ^34^, showing Siglec-15 binding increase (ΔMFI) upon mucin expression in HEK^WT^ or STn cells (HEK^KO:COSMC;KI:ST6GALNAC1^). **(e)** ELISA analysis of Siglec-15 binding to select purified recombinant mucin reporters with sulfo-dST O-glycans; values represent mean ± SEM from n = 4.

The binding of Siglec-7-Fc to HEK293 cells expressing CHST1 almost reached signal saturation at 1 µg/mL (**Fig. 2b**), masking any potential differences in binding upon expression of mucin reporters. Therefore, we first titrated the Siglec-7-Fc concentration and found that 250 ng/mL precomplexed Siglec-7-Fc provided low detectable binding (MFI > 300-fold above background) to HEK293 cells with 6-*O*-sulfation (HEK^KO:GCNT1;KI:CHST1^) and without expression of mucins (**Fig. 2b**). Using this assay condition, Siglec-7-Fc binding was found to be highly dependent on the mucin reporters expressed in these sulfo-dST glycoengineered cells (HEK^KO:GCNT1;KI:CHST1^). In contrast, binding was largely unaffected when the same mucin reporters were expressed in cells engineered to produce sulfo-mSTa O-glycans without α2-6-sialylation (HEK^KO:GCNT1/ST6GALNAC2-4;KI:CHST1^), with weak binding only induced by MUC3A, MUC17, and MUC22 in these cells (**Fig. 3b**). Several mucin reporters induced a strong increase in binding, including those designed from mucin-like domains of the previously reported Siglec-7 ligands CD43 and PSGL-1 ^40,61^, but also previously unknown ligands such as MAdCAM1, which induced the highest binding increase. In contrast, classical mucins with high density (e.g., MUC5Ac and MUC2) or low density (e.g., MUC1 and MUC20) of O-glycans did not enhance binding (**Fig. 3b**). The differential binding of Siglec-7-Fc to mucin-like domains was reproduced with select recombinant secreted mucin reporters by ELISA (**Fig. 3c**). Interestingly, a different set of mucins induced Siglec-7 binding when expressed in cells engineered with STn O-glycans (**Fig. 3b, inset**, data generated in our previous study ^34^). For example, MUC2 and MUC5Ac (with high-density of O-glycans) strongly induced Siglec-7-Fc binding when expressed in STn cells, while we did not observe any binding increase upon expression in cells engineered with sulfo-dST O-glycans (HEK^KO:GCNT1;KI:CHST1^).

Siglec-15 binds weakly to STn on glycoengineered HEK293 cells and binding is markedly augmented by expression of most of the mucin reporters tested ^34^. However, when CHST1 sulfation is introduced to HEK293 cells that do not express STn O-glycans or mucin reporters, this also leads to marked Siglec-15-Fc binding in an O-glycan dependent manner ^34^. To probe into this further, we first titrated the Siglec-15-Fc concentration finding that 1 µg/mL precomplexed Siglec-15-Fc provided low detectable binding (MFI > 60-fold above background) to HEK cells with CHST1 sulfation (HEK^KO:GCNT1;KI:CHST1^) and without expression of mucins (**Fig. 2b**). Using this assay condition, Siglec-15-Fc binding was highly dependent on the expression of mucin reporters in cells engineered to produce sulfated dST (HEK^KO:GCNT1;KI:CHST1^), as well as sulfated mSTa O-glycans (HEK^KO:GCNT1/ST6GALNAC2-4;KI:CHST1^) without α2-6-sialylation (**Fig. 3d**). Similar to Siglec-7, we observed a high increase in Siglec-15-Fc binding with MAdCAM1, CD43 and PSGL-1 mucin reporters, and this binding could be reproduced with the purified recombinant secreted sulfated mucin reporters by ELISA (**Fig. 3e**). Of note, some reporters (i.e., MUC3A, MUC6, CD43) showed higher Siglec-15-Fc binding when expressed on sulfo-mSTa cells compared to sulfo-dST. We also observed that Siglec-15-Fc bound to a different set of mucin reporters when expressed on STn cells, including MUC2 and MUC5Ac (**Fig. 3d**) ^34^.

Given the important effects of CHST1 6-*O*-sulfation, we further explored other sulfotransferases, including the 6-*O*-sulfotransferases CHST2-6 and the 3-*O*-sufotransferases GAL3ST2/4 (**Supplementary Fig. 2**). Siglec-7-Fc binding to mucin reporters was also enhanced to variable extents by other sulfotransferases except for the 6-*O*-sulfotransferase CHST3 and the 3-*O*-sulfotransferase GAL3ST2 (**Supplementary Fig. 2a**), similar to Siglec-7-Fc binding data previously obtained in the absence of mucins ^34,37^. The enhanced binding was preferentially observed with the MAdCAM1 and PSGL-1 reporters, while e.g. the MUC5Ac reporter did not induce binding. Notably, the 3-*O*-sulfotransferase GAL3ST4 also enhanced binding of Siglec-7-Fc but only in combination with expression of MAdCAM1 and PSGL-1 (weakly MUC13), and not CD43, which has previously been identified as a Siglec-7 ligand ^34,40^. Siglec-15-Fc binding to most mucin reporters was enhanced almost exclusively by introduction of CHST1-mediated 6-*O*-sulfation (**Supplementary Fig. 2b**). In contrast, introduction of other sulfotransferases only enhanced Siglec-15-Fc binding in combination with the PSGL-1 reporter, but not with any of the other reporters.

These results clearly point to a role of sulfate groups in Siglec-7/15 binding and further indicate that the position of such sulfate groups on Gal or likely GlcNAc (of core2 O-glycans) may not be decisive for the observed enhanced binding, while the carrier O-glycoprotein and the ST O-glycan is critical for binding. Note though that due to analytical challenges we have not addressed the structures of the sulfo-ST O-glycans involved in the binding epitopes, and it is possible that the enhanced binding by sulfation is not mediated by direct involvement in the binding epitope, but perhaps through indirect effects that expose the binding epitope.

### Select O-glycoproteins with clustered O-glycan motifs guide Siglec-7 and −15 binding

We first employed our recently developed Glycocarrier platform to probe simple O-glycan cluster motifs comprised of 1-6 adjacent O-glycans on Ser/Thr residues (**Fig. 4a**). Glycocarriers are designed to resemble mucin reporters (**Fig. 3a**), but the mucin-like glycomodules (around 200 amino acids) contain identical or different short O-glycosylation sequence motifs (10-20 amino acids) in tandem repeats interspaced by trypsin cleavage sites (**Supplementary Table 2**) ^62^. As both Siglec-7-Fc and Siglec-15-Fc bind to mucins displayed on cells with capacity for sulfo-dST (HEK^KO:GCNT1;KI:CHST1^) (**Fig. 3**), we first probed Siglec binding to Glycocarriers expressed in this cell line. Surprisingly, we did not observe enhanced binding of Siglec-7- or Siglec-15-Fc, while expression of the MAdCAM1 reporter clearly enhanced binding (**Fig. 4a**). We also expressed the Glycocarriers in STn engineered cells (HEK^KO:C1GALT1;KI:ST6GALNAC1^) without observing any binding (**Supplementary Fig. 3a**). These findings suggest that simple O-glycan cluster motifs are not sufficient to guide Siglec binding.

**Figure 4.**
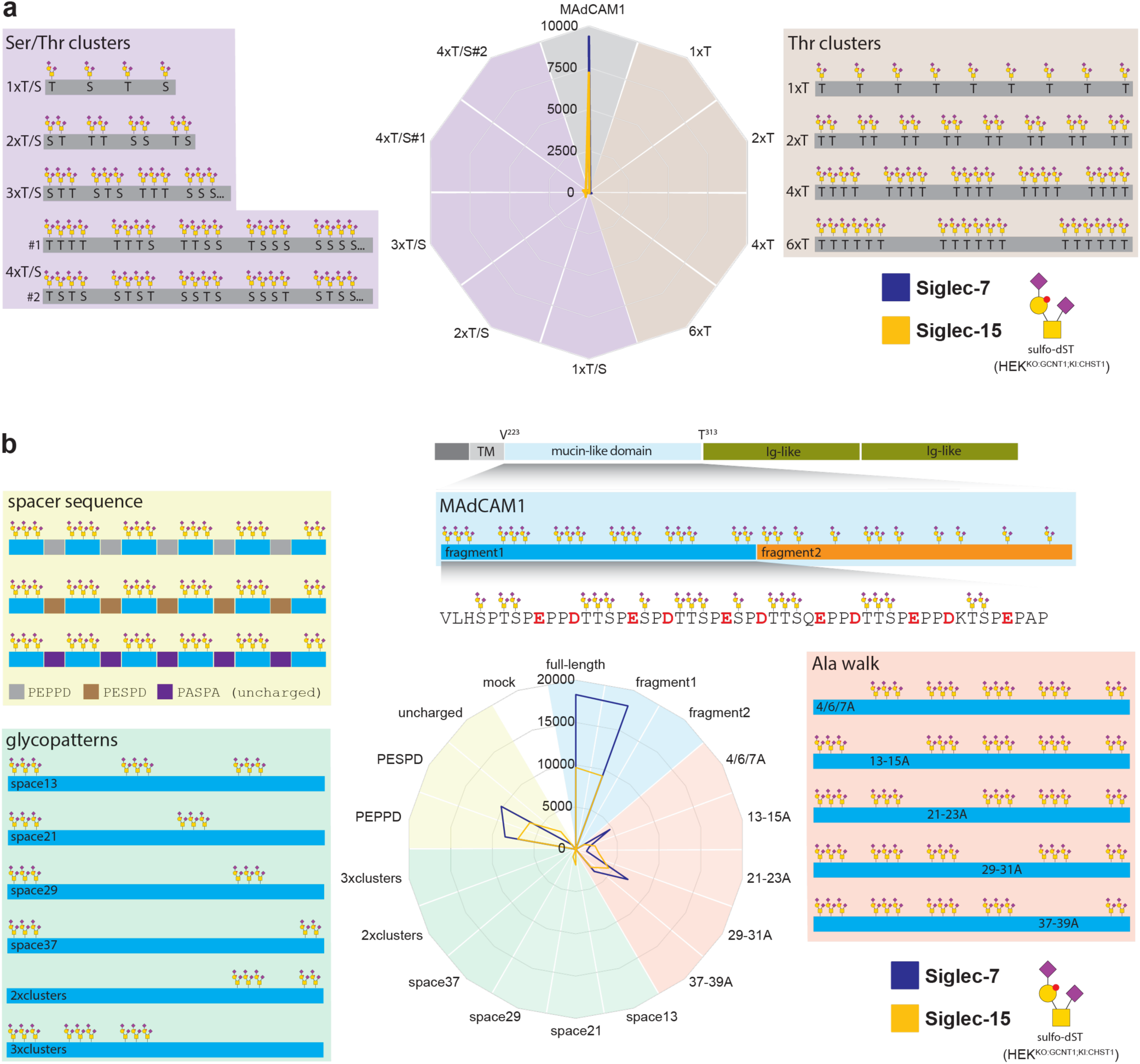
Flow cytometry analysis of putative O-glycan cluster motifs for Siglec-7/15 binding by Glycocarriers and mutant MAdCAM1 reporters. **(a**) Binding of Siglec-7 (blue) and Siglec-15 (yellow) to a library of rationally designed O-Glycocarriers displayed on cells engineered to produce sulfo-ST O-glycans (HEK^KO:GCNT1;KI:CHST1^) as indicated. Color code of radar plots: Brown, clusters of 1-5 Thr repeats; Purple, clusters of 1-4 Ser/Thr repeats. Representative MFI values from two independent experiments. Glycocarrier sequences are listed in **Supplementary Table 2.** (**b**) Binding of Siglec-7 and Siglec-15 to a library of MAdCAM1-based reporters displayed on sulfo-ST cells (HEK^KO:GCNT1;KI:CHST1^). Color code of radar plots: Blue, MAdCAM1 (Figure 4 legend continued) fragments; Red, alanine walk mutants (removal of individual TTS repeats); Green, O-glycan cluster mutants (deletion of TTS repeats); Yellow, Spacer mutants. Mock-transfected cells were subjected to the same transfection protocol but leaving out DNA. Representative MFI values from four independent experiments. Cell surface expression of the Glycocarriers and MAdCAM1 reporters was verified with an anti-FLAG mAb. Sequences of MAdCAM1-based reporters are listed in **Supplementary Table 3**.

This prompted us to use the MAdCAM1 reporter, which showed the highest Siglec-7-Fc binding in our mucin display screening (**Fig. 3b**), as a model O-glycoprotein to narrow down features required for binding of Siglec-7/15 (**Fig. 4b, Supplementary Table 3**). The MAdCAM1 O-glycomodule contains different N- and C-terminal O-glycan motifs, so we first tested which of these fragments guided Siglec binding when expressed in cells engineered to produce sulfo-dST (**Fig. 4b**) and dST (**Supplementary Fig. 3b**). This revealed that the N-terminal region (fragment1) of the MAdCAM1 O-glycomodule with repeated TTS triad O-glycan motifs exclusively directed binding of both Siglec-7- and Siglec-15-Fc (**Fig. 4b** and **S2b**). Similar binding patterns were found when the MAdCAM1 reporters were expressed in dST cells with or without CHST1-mediated 6-*O*-sulfation, albeit with significantly lower intensity in the absence of sulfation (**Supplementary Fig. 3b**).

We proceeded with an “alanine walk” strategy where individual TTS motifs were mutated to AAA in the MAdCAM1 fragment1 reporter (**Supplementary Table 3**); however, removal of any of the single TTS repeats resulted in significantly reduced Siglec-7/15-Fc binding (**Fig. 4b**). We also created a library of different patterning of the TTS repeats where several adjacent or distanced TTS repeats were mutated to AAA. Interestingly, a complete loss of binding was observed when removing more than one TTS repeat, regardless of whether the mutated TTS repeats were adjacent or spaced further apart from each other (**Fig. 4b**).

Finally, we rationally designed reporters with reduced complexity based on the MAdCAM1 fragment1 sequence with homogenous spacer regions between the TTS clusters of PEPPD or PESPD, respectively, or by mutating all charged residues in the spacer region to alanine (**Supplementary Table 3**). Siglec-7-Fc and Siglec-15-Fc consistently recognized the PEPPD and PESPD reporters, albeit with lower binding intensity compared to the natural fragment1 sequence. In contrast, complete loss of binding was observed upon removal of charged residues from the spacer region (**Fig. 4b**).

These results suggest that the repeated O-glycan clusters as well as the acidic amino acid residues in the spacer sequences in-between these clusters are important for Siglec-7/15 binding. While the repeat motifs likely support multivalent interactions with the oligomeric Siglecs, the need for acidic residues in-between these repeated O-glycan clusters is less apparent, and we therefore hypothesized that these residues play a role in directing CHST1 sulfation.

### CHST1-mediated 6-O-sulfation is sequence-dependent and a key factor for optimal Siglec-7 and −15 binding

Current knowledge of enzymatic sulfation of O-glycans is still limited. We previously introduced CHST1/3 6-*O*-sulfotransferases to HEK293^KO:COSMC^ cells with truncated Tn O-glycans and were unable to demonstrate significant 6-*O*-sulfation of a secreted MUC1 reporter; however, we did not explore 6-*O*-sulfation of extended ST/T O-glycans ^63^. Here, we first performed sulfoglycomics on HEK^KO:GCNT1;KI:CHST1^ cell lysates and detected both sulfo-dST and sulfo-mST O-glycans in combination with non-sulfated core1 O-glycans (**Supplementary Fig. 4b**), which indicated that sulfation on O-glycans was present but incomplete. Note that while our sulfoglycomics analysis indicated incomplete sulfation, it is well established that quantification of sulfated and non-sulfated O-glycans is challenging ^64^.

We next focused on sulfation of O-glycans on the reporters expressed as secreted fusion proteins in HEK^KO:GCNT1;KI:CHST1^ cells (**Supplementary Fig. 4c**). Surprisingly, while we identified sulfo-dST and sulfo-mST O-glycans together with the non-sulfated species on the MAdCAM1 (full-length and fragment1) and CD43 reporters, we could not identify sulfated O-glycans on the MUC5AC reporter. The MUC5AC reporter sequence stands out by not containing acidic Glu/Asp residues (**Supplementary Table 1**), and removal of Glu/Asp residues in the MAdCAM1 reporter abrogated Siglec-7-Fc binding (**Fig. 4b**), suggesting that these negatively charged residues in fact could be contributing to efficiency of the sulfation by CHST1. To explore this, we analyzed different isolated MAdCAM1 reporter variants produced in HEK^KO:GCNT1;KI:CHST1^ cells by sulfoglycomics (**Supplementary Fig. 5**). This revealed the presence of sulfated dST and mST O-glycans in all the tested fragment1 reporters (29-31A, space13, uncharged) as well as the fragment2 reporter (**Supplementary Fig. 5**). Thus, CHST1 sulfation does not appear to depend on particular features of the MAdCAM1 fragment1 sequence.

To further estimate the sulfation efficiency on these reporters, we next characterized a subset of MAdCAM1 reporters by ELISA (**Supplementary Fig. 6**). We used the PNA lectin recognizing core1 (Galβ1-3GalNAcα/β), but not sulfo-core1 ^63^, to probe for potential masking of core1 by sulfation, which requires prior removal of sialic acids by neuraminidase treatment to expose core1 underlying ST and sulfo-ST O-glycans. Interestingly, PNA binding to neuraminidase-treated MAdCAM1 reporters produced in HEK cells with CHST1 6-O-sulfation capacity (HEK^KO:GCNT1;KI:CHST1^) was almost completely abrogated compared to reporters produced in cells without CHST1 expression (HEK^KO:GCNT1^), with the notable exceptions of the MAdCAM1 full-length, uncharged fragment1, and fragment2 reporters where the binding of PNA was largely unaffected (**Supplementary Fig. 6**). This suggests that fragment1 is more efficiently sulfated than fragment2, more than predicted from our glycomics analysis, which revealed only minor sulfated and mainly non-sulfated ST O-glycans on both fragments. Importantly, CHST1 sulfation was not affected by most of the sequence modifications introduced to fragment1 MAdCAM1, including changes in spacing (“space13”) and number (“29-31A”) of the O-glycan clusters, as PNA binding was almost completely abrogated. In contrast, removal of the charged residues in between the O-glycan clusters (“uncharged”) left PNA binding relatively unperturbed. This suggests that the sulfation efficiency as evaluated by blocking of PNA binding is lower in the uncharged MAdCAM1 reporter.

Taken together, these results indicate that optimal binding of precomplexed Siglec-7-Fc to MAdCAM1 requires multiple TTS repeats and sulfation of O-glycans. Interestingly, Siglec-15 appeared to share the same MAdCAM1 binding requirements, albeit with weaker intensity. Our finding that a large fragment of the O-glycodomain of MAdCAM1 with multiple TTS repeats was needed for binding further suggests that a multivalent interaction drives Siglec binding. This would agree with previous studies demonstrating that precomplexing of Siglec-Fc chimera is important to detect binding with Siglecs ^47,65^, and that the repeated O-glycan clusters in the select MAdCAM1, CD43, and PSGL-1 ligands are appropriately spaced to support such multivalent interactions.

### MAdCAM1 reporters support binding to Siglec-7 installed on CHO cells

The need for precomplexing of Siglec-Fc chimeras for detectable binding to ligands poses questions of the binding properties of Siglecs when they are naturally presented on cells ^47,66,67^ (**Fig. 1**). We therefore explored the binding of purified recombinant secreted mucin reporters to CHO cells stably expressing Siglec-7 (CHO^Sig-7WT^), with cells expressing a Siglec-7 R124A inactive mutant as control (CHO^Sig-7R124A^) (**Fig. 5a**). We initially tested purified recombinant MAdCAM1 reporters using their N-terminal GFP for binding detection, and found only weak shifts in fluorescence intensities, so we employed AF488-conjugated MAdCAM1 reporters for all subsequent cell binding assays.

**Figure 5.**
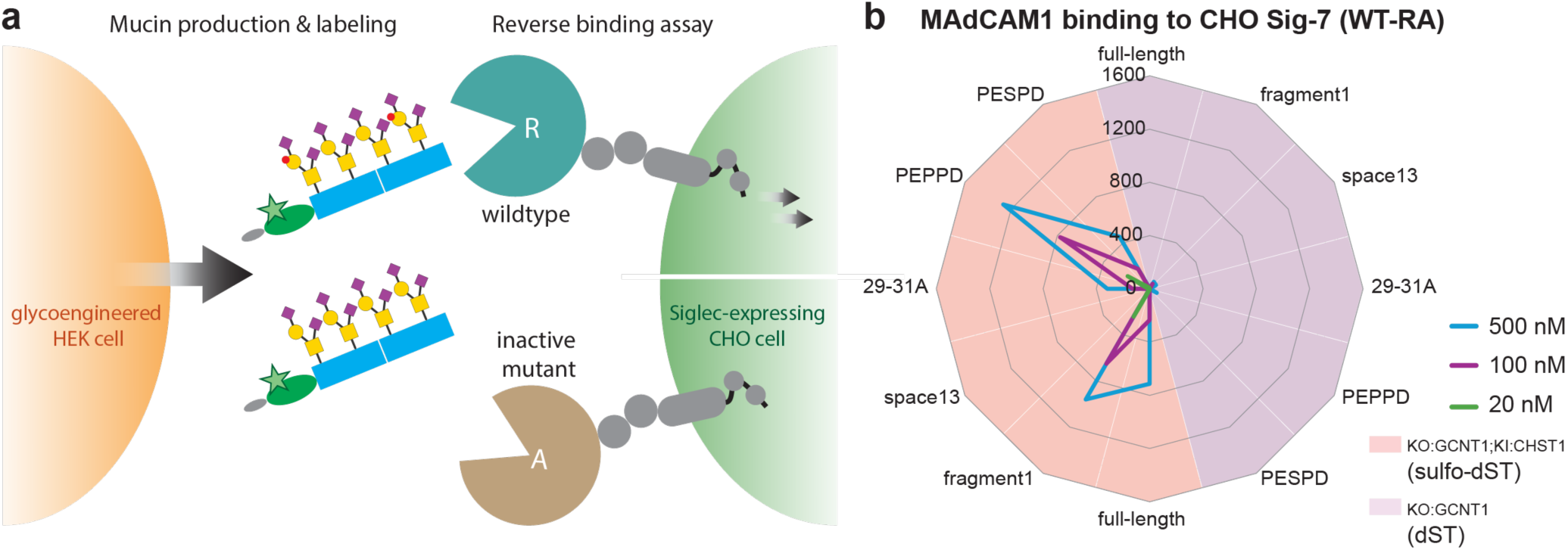
Flow cytometry analysis of binding to Siglec-7 installed on CHO cells. (**a**) Workflow of the reverse Siglec binding assay. Mucins secreted by glycoengineered HEK cells were purified and labelled with a fluorophore. Binding of labelled reporters to CHO^Sig-7WT^ and CHO^Sig-7R124A^ cells was then probed. (**b**) Binding of non-sulfated (purple) vs. sulfated (salmon) MAdCAM1-based reporters to CHO^Sig-7^ cells at different concentrations. Values represent MFI obtained by subtracting MFI (CHO^Sig-7R124A^) from MFI (CHO^Sig-7WT^), mean from two independent experiments.

We probed the dST and sulfo-dST glycoforms of AF488-conjugated MAdCAM1 reporters with CHO cells expressing Siglec-7. Importantly, we found robust binding only of the sulfo-dST glycoform and not the dST glycoform (**Fig. 5b**), while neither of these glycoforms bound to the Siglec-7 mutant-displaying CHO^Sig-7R124A^ cells (**Supplementary Fig. 7**). Similar to our studies of binding with precomplexed Siglec-7-Fc to MAdCAM1 reporters displayed on cells (**Fig. 4b**), the isolated AF488-conjugated MAdCAM1 variants showed that binding was mediated only by the N-terminal region (fragment1) with preference for the PEPPD spacer sequence in between TTS O-glycan clusters, and removal of a single (29-31A) or multiple TTS motifs (space13) resulted in a reduction or loss of binding, respectively. Note that binding of the sulfo-dST MAdCAM1 reporter (full-length and fragment1) was detectable down to a concentration of 20 nM (**Fig. 5b**). These results suggest that the sulfo-dST O-glycan clusters (repeated TTS motifs interspaced with short sequences containing one acidic Glu/Asp) in the N-terminal region of MAdCAM1 provide an optimal binding epitope for Siglec-7.

Next, we tested select reporters for which we already showed that precomplexed Siglec-Fc binds (CD43, PSGL-1) or does not bind (MUC5AC) (**Fig. 3b-e**), and the purified AF488-conjugated reporters revealed the same binding specificities with CHO^Sig-7^ cells (**Supplementary Fig. 8**). In striking contrast to our observations with precomplexed Siglec-7-Fc, no binding was observed with the non-sulfated variants of MAdCAM1, CD43, or PSGL-1 **(Fig. 5, Supplementary Fig. 8)**, clearly demonstrating that CHST1-mediated 6-*O*-sulfation is a determining factor in reaching sufficient affinity to engage Siglec-7 displayed on cells. To explore potential *cis*-masking of Siglec-7 on CHO^Sig-7^ cells, we generated a CHO^Sig-7;KO:Cosmc^ with KO of *Cosmc* to eliminate elongation and sialylation of O-glycans, as the main *cis*-ligands for Siglec-7 are sialo-O-glycans ^38^. CHO^Sig-7;KO:Cosmc^ and CHO^Sig-7^ cells showed essentially the same binding profiles with the mucin reporters, suggesting that *cis*-masking is not relevant in this cell system (**Supplementary Fig. 9**).

To evaluate selectivity of sulfo-dST MAdCAM1 for Siglec-7, we screened CHO cells installed with Siglec-3, −8, or −15 (CHO^Sig-3^, CHO^Sig-8^, CHO^Sig-15^) (**Supplementary Fig. 7**). We previously found that binding of these Siglecs was highly sensitive to CHST1-mediated 6-*O*-sulfation with precomplexed Siglecs using the cell-based mucin display ^34^. Probing these cell lines revealed that Siglec-7 was the only receptor for sulfated MAdCAM1 (and fragment1).

Interestingly, cells expressing Siglec-15 did not exhibit appreciable binding (**Supplementary Fig. 7**), in contrast to what was observed with precomplexed Siglec-15-Fc and MAdCAM1 displayed on HEK^KO:GCNT1;KI:CHST1^ cells (**Fig. 4**). The reason for this is unclear at present and may relate to poor and unstable Siglec-15 expression levels on CHO cells or to lower accessibility of the glycan-binding site as it is positioned closer to the membrane compared to Siglec-7 ^2^. However, we note that Siglec-15 in general showed weaker binding than Siglec-7 to glycoengineered cells in titration experiments (**Fig. 2b**), and that we used four times higher concentration of Siglec-15-Fc to achieve binding intensities similar to that of Siglec-7-Fc (**Fig. 2b, Fig. 3b/d**).

Two recent studies reported ligand-induced clustering to be important for the function of Siglec-7 ^29,50^, and we therefore examined MAdCAM1-induced oligomerization and internalization of Siglec-7 with CHO^Sig-7^ cells by confocal microscopy. Indeed, sulfated MAdCAM1 showed binding to CHO^Sig-7WT^ as distinct puncta, indicative of clustering of Siglec-7 on the cell surface **(Supplementary Fig. 10)**. To evaluate uptake of the sulfated MAdCAM1 reporter by CHO^Sig-7^, we monitored specific uptake at 37°C in a time course, finding internalization after 30 min (**Supplementary Fig. 10**), while no binding or uptake was observed with CHO^Sig-7R124A^ cells. Uptake of sulfated MAdCAM1 thus occurs on a similar time scale compared to previous studies using liposomes or antibodies to induce internalization of Siglec-3 and −9 ^68,69^.

### Sulfo-dST O-glycosylated MAdCAM1 targets primary human monocytes that express Siglec-7

To further explore the selectivity of sulfo-dST MAdCAM1 for Siglec-7, we probed binding of the fluorophore-tagged MAdCAM1 glycoforms to isolated primary immune cells (peripheral blood mononuclear cells, PBMCs) from healthy human donors (**Fig. 6a**). We found that sulfo-dST MAdCAM1 selectively binds to a distinct subset of cells (∼13%) in all six tested healthy donor PBMC populations, while the non-sulfated dST glycoform of MAdCAM1 did not show binding (**Fig. 6b**). We confirmed that the sulfo-dST MAdCAM1-binding immune cells expressed Siglec-7 by anti-Siglec-7 antibody co-staining, with >80% of Siglec-7^+^ cells positive for binding of sulfo-dST MAdCAM1.

**Figure 6.**
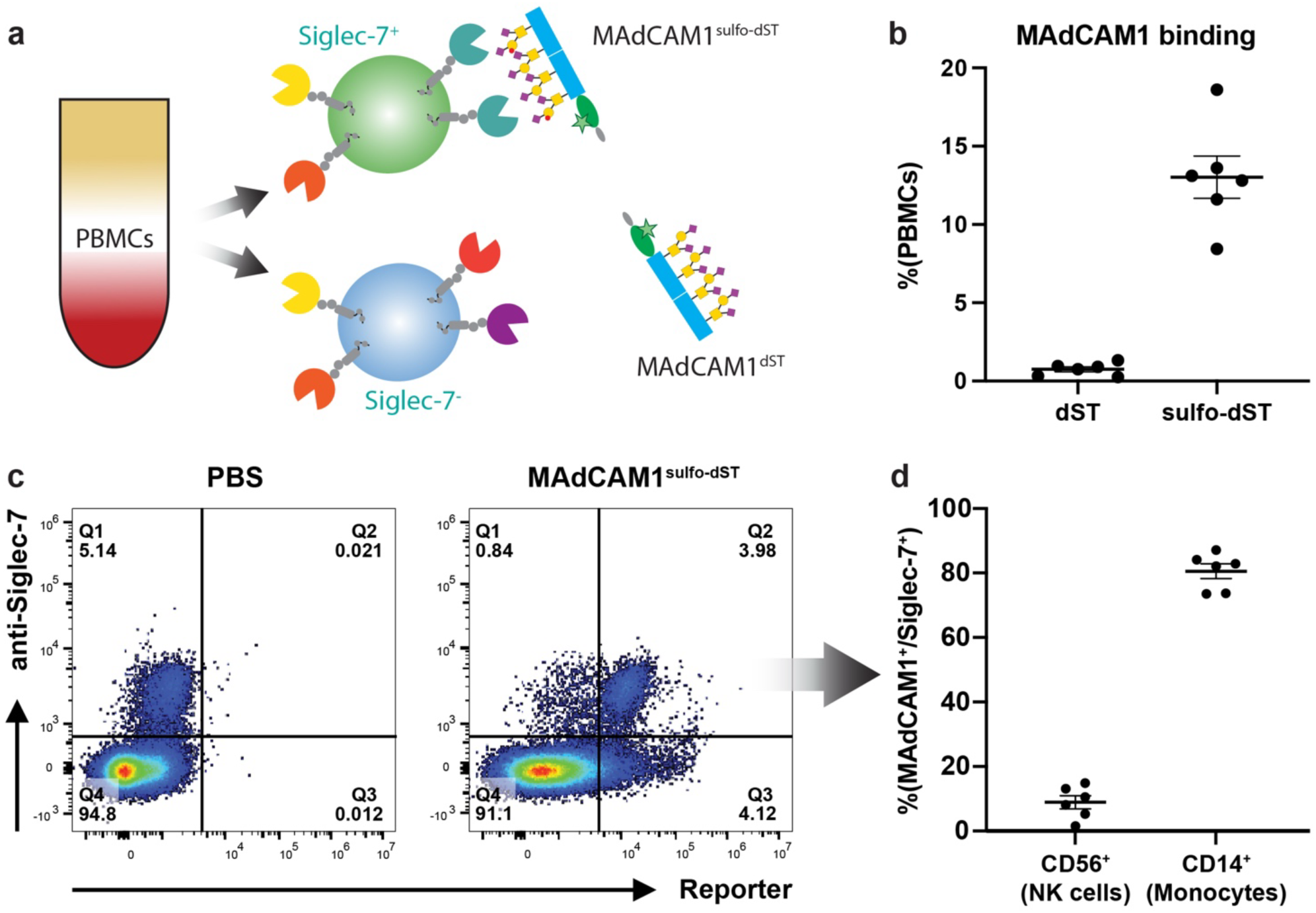
Flow cytometry analysis of MAdCAM1 binding to PBMCs. (**a**) Graphic depiction of the immune cell binding assay. PBMCs were isolated by density gradient centrifugation and incubated with Alexa488-labelled MAdCAM1 glycoforms as indicated. (**b**) Binding (% of total PBMCs) of MAdCAM1 glycoforms in six healthy human donors (mean ± SEM). (**c**) Staining with anti-Siglec-7 antibody ± MAdCAM1^sulfo-dST^. Note the MAdCAM1^+^/Siglec-7^+^ population in Q2 used for further analysis in (d). (**d**) Co-staining with anti-CD56, and anti-CD14 antibodies to identify NK cells and monocytes, respectively, in the MAdCAM1^+^/Siglec-7^+^ population. The percentage of CD56^+^ or CD14^+^ cells in that population is displayed for six healthy human donors (mean ± SEM).

Interestingly, despite similar Siglec-7 expression levels on Siglec-7^+^ monocytes (CD14^+^) and NK cells (CD56^+^) (**Supplementary Fig. 11**), we detected strong binding of sulfo-dST MAdCAM1 to essentially all Siglec-7^+^ monocytes, but hardly any binding to Siglec-7^+^ NK cells (**Fig. 6c-d, Supplementary Fig. 11**). The PBMCs contained two-fold higher numbers of monocytes (∼20%) than NK cells (∼10%) (**Supplementary Fig. 11**), leaving no simple explanation for the observed selective binding of sulfo-dST MAdCAM1 to monocytes.

Siglecs are inhibited by *cis-*ligands on the same immune cells ^66^, and neuraminidase pretreatment of NK cells to remove sialic acids resulted in unmasked binding of synthetic Neu5Ac(α2-8)Neu5Ac-polyacrylamide (Sia-PAA) probes ^57^. In the same study, the authors reported that human Siglec-7 expressed in CHO cells did not experience *cis*-inhibition, in agreement with our results (**Supplementary Fig. 9**). We therefore considered that different levels of *cis-*inhibition in monocytes and NK cells could explain our finding that sulfo-dST MAdCAM1 bound well to monocytes but not NK cells ^2,57^. However, neuraminidase pretreatment of PBMCs to remove sialic acids did not significantly change the selectivity of the sulfo-dST MAdCAM1 reporter for monocytes compared to NK cells (**Supplementary Fig. 11)**. Of note, the monocyte population rather homogenously expressed Siglec-7, whereas Siglec-7 expression on NK cells was more heterogeneous (ranging from 20-90% expression between donors). These results suggest that sulfo-dST MAdCAM1 serves as a ligand for Siglec-7 on monocytes, and further studies are needed to clarify the different binding properties of Siglec-7 on NK cells.

## DISCUSSION

Our study provides evidence for the existence of contextual O-glycoprotein ligand motifs for Siglec-7 and −15, and it illustrates how such complex motifs can be unraveled using recombinant O-glycodomain reporters designed from native human glycoproteins and displayed or produced in genetically glycoengineered cells. Molecular structural studies will be needed to define the exact binding motifs and mode of recognition, and our study also does not address putative signaling and functional effects following binding of the reporters to Siglecs. However, our results support the original hypothesis that the limited set of simple sialic acid capped glycan epitopes with widespread expression in human tissues cannot provide sufficient ligand diversity to orchestrate select regulation of endogenous lectin receptors like Siglecs and Selectins ^1,25–27^. We found intriguing evidence that Siglec-7/15 recognize shared clusters of sulfated O-glycans on select O-glycoproteins like MAdCAM1, CD43, and PSGL-1, although with a subtle difference in their dependence on α2-6 sialic acid on the inner GalNAc residue in core1 O-glycans (dST vs. mST sulfo-O-glycans). Moreover, the sulfation of O-glycans on these reporters by CHST1 seems to (at least partly) determine their role as ligands for Siglec-7/15. Finally, the identified O-glycoprotein ligands all contained repeated motifs of O-glycan clusters pointing to the involvement of multivalent interactions with oligomeric Siglec assemblies ^29,50,51^. These results suggest that the engagement of Siglec-7/15 receptors on immune cells involves the concerted expression of particular O-glycoproteins and select sialyl- and sulfotransferase isoenzymes to produce and display repeated sulfo-sialo O-glycan motifs.

Our study highlights the influence of assay design in binding studies with GBPs like Siglecs and illuminates the need to look beyond simple glycan epitopes and continue searching for contextual ligands with better and select binding. While binding studies with simple glycan epitopes clearly are informative and instructive, we found that only the best ligands identified in direct binding assays with precomplexed Siglec-7-Fc ultimately resulted in detectable binding to Siglec-7 naturally expressed on cells (**Fig. 6**). Thus, binding studies with simple glycans may not be sufficient to appreciate how GBPs achieve selectivity through recognition of contextual presentation of glycans on proteins and arrangements for multivalent interactions. The cell-based glycan array platform offers opportunities to discover and dissect such contextual glycoprotein epitopes, because glycoproteins and O-glycodomain reporters can be expressed and displayed with custom-designed, rather homogenous glycans in the natural context of the protein backbone ^70,71^. Moreover, this platform enables reverse assay designs where recombinant secreted O-glycoprotein reporters are isolated and tagged to probe binding and uptake by endogenously expressed GBPs on cells in their natural oligomeric state and other assemblies at the cell membrane ^31,62^.

Our glycan array was built in HEK293 cells, and these do not endogenously express mucins, including the mucin-like O-glycoprotein targets for Siglec-7/15 we identified here. Our cell-based array strategy thus relies on prior knowledge of candidate glycoprotein targets and the recombinant expression of these in appropriately glycoengineered cells, such as the mucin-like reporters employed in this study. The cell-based array concept is, however, widely applicable to cell lines that may endogenously express relevant target O-glycoproteins and are amenable to genetic glycoengineering. This was demonstrated with the human promonocytic leukemia cell line U937 where overexpression of CHST1 greatly enhanced binding of Siglec-7-Fc in an O-glycan-dependent manner ^37^. We note that U937 cells endogenously express two identified targets for Siglec-7, PSGL-1 and CD43 ^73^, and a dissection strategy with knockout of these genes in U937 cells may be employed to validate Siglec binding to these endogenous O-glycoproteins. More generally, identification of specific glycoprotein ligands with contextual Siglec binding motifs may include genome-wide CRISPR knockout screenings as well as affinity isolation of targets. These strategies were in fact previously used to identify both CD43 and PSGL-1 as Siglec-7 ligands ^40,41,61^. Different and complementary approaches may therefore be adopted to identify glycoproteins with contextual glycan binding motifs that serve as natural ligands for Siglecs.

We also demonstrate the importance of employing the “reverse binding assay” format with isolated tagged glycoproteins serving as bait for GBPs expressed on cells. For this, we used recombinant secreted reporters designed to contain extended O-glycodomains from mucin-like O-glycoproteins to capture the full context of O-glycan clusters and repeated motifs found in these. Currently, the cell-based array platform is the only practical way to produce such large and complex biomolecules, but it also poses challenges when it comes to detailed structural analysis and the complete removal of impurities that may affect cell behavior. The latter is exacerbated by the high number of negative charges associated with sialic acids and sulfate groups decorating the densely spaced O-glycans in mucins. This is particularly apparent when removing the N-terminal GFP modules from our mucin reporters by TEV cleavage, where the heavily charged mucin reporters precipitate in neutral buffers. In this study, we therefore only used the secreted isolated reporters with GFP for binding studies at cold temperature.

We found that only the sulfo-dST O-glycoform of select O-glycoproteins supported strong binding to Siglec-7 expressing CHO cells (**Fig. 5**) and human monocytes (**Fig. 6 and Supplementary Fig. 11**) – in striking contrast to precomplexed Siglec-7-Fc and Siglec-15-Fc, which show binding to a wide range of sialylated O-glycoproteins ^34,40^. Interestingly, we found only weak/no binding to human NK cells, and, surprisingly, this lack of binding was not due to *cis*-inhibition as removing sialic acids by neuraminidase pretreatment did not induce binding (**Supplementary Fig. 11**). Currently, we are unable to explain this finding. However, differential binding to Siglec-7 displayed on NK cells and monocytes was also previously found ^57^. In this study, synthetic Sia-PAA was found to selectively bind CD56^+^ NK cells only after treatment with neuraminidase to remove *cis*-ligand blocking; however, the Sia-PAA did not bind to other CD56^-^ cells in the PBMC population, including monocytes. Thus, it appears that the synthetic Sia-PAA ligands behaves differently from our O-glycoprotein reporters in binding to Siglec-7 on immune cells, and further studies are clearly needed to resolve these differences. An obvious limitation stems from the use of ligands in solution for binding to Siglec-expressing cells. While our secreted reporters have the unique advantage of enabling the reverse binding assay format, we still need to develop technologies for dissecting O-glycoprotein-Siglec interactions in a cell-cell format. In our current cell-based array platform, one major roadblock for cell-cell assays is that the engineered O-glycans displayed on the glycoprotein ligands of interest will inherently also be found on endogenous glycoproteins. This is the reason for the “background” subtraction needed to study Siglec binding to select mucin reporters displayed on glycoengineered cells (such as in Fig. 3).

Surprisingly, the same O-glycoprotein reporters that served as efficient ligands for Siglec-7 (MAdCAM1 fragment1, CD43, and PSGL-1) were also those that seemed to acquire higher 6-*O*-sulfation levels (**Supplementary Fig. 4-5**). Thus, our study points to select sulfo-sialo O-glycoproteins as the endogenous ligands and thereby to sulfotransferases like CHST1 as key regulatory enzymes for the formation of Siglec-7 ligands. However, how these O-glycoproteins are selected for CHST1 sulfation to become Siglec-7 ligands is unclear. We found clustered O-glycan motifs interspaced by negatively charged Glu/Asp residues (**Supplementary Fig. 8**) to be a common feature of the identified Siglec-7 ligands (**Fig. 4**), although deleting these acidic residues only reduced but did not abrogate sulfation (**Supplementary Fig. 4-5**). Knowledge of O-glycan sulfation and the substrate specificity of the enzymes involved is still very limited, partly due to analytic challenges with identifying and quantifying sulfated O-glycans ^74^, and partly because the properties of the many different sulfotransferases and their roles in sulfating O-glycans have not been studied ^63^.

Interestingly, previous screenings for Siglec-7 ligands frequently employed cell lines with low or no expression of CHST1, such as the K562 cell line used for identification of CD43 as a Siglec-7 ligand ^35,40,42^. Further studies are thus clearly needed, and CHST1 is a particularly interesting candidate since this sulfotransferase is overexpressed in a variety of cancers and its expression correlates with poor patient outcome ^37^.

The fine molecular nature of the contextual sulfo-ST O-glycoprotein binding epitopes for Siglec-7 is challenging to narrow down and will require further studies. The three identified O-glycoprotein targets MAdCAM1, CD43, and PSGL-1 highlight that clustered sulfo-sialo O-glycans as well as repeats of these clusters are important for Siglec binding (**Supplementary Fig. 8**). The V-set Ig-like binding domain of Siglecs with a conserved arginine residue (Arg124 in Siglec-7) accommodates binding to a single sialic acid residue ^54,55^, but additional noncanonical binding sites for sialic acids ^60^ and glycolipids ^67^ may involve another conserved Arg residue (Arg94 in Siglec-7). Siglec-7 is reported to bind disialogangliosides ^3,23,56–58^; however, these are likely not displayed on HEK293 cells since loss of elaborate O-glycans (KO COSMC/C1GALT1) fully abrogates Siglec-7 binding ^34^. An NMR study of Siglec-8 binding to synthetic 6-sulfo-sialyl-Lewis-X glycans found specific interactions between the C’C loop and the sulfate group ^75^, and it is conceivable that Siglec-7 also interacts directly with sulfate groups in sulfo-ST O-glycoproteins. Here, we tried to dissect the contextual O-glycan epitopes with the MAdCAM1 reporter by truncations, mutations, and pattern modifications of the O-glycan clusters. Our results point towards a contextual epitope comprised of sulfo-ST structures in clusters of 2-3 adjacent O-glycans and with a requirement for repeated cluster epitopes, likely spaced to support multivalent high-avidity interactions by the natural oligomeric state of Siglec-7 (**Fig. 1**). Importantly, the dissection of complex binding properties as found for these Siglecs relies on faithful glycosylation and sulfation of the glycoprotein reporters displayed in glycoengineered cells. While our previous studies and data presented here provide support for faithful O-glycosylation of the reporters, we have not in detail characterized the occupancy of O-glycans and the structures of O-glycans at each position of the glycosites in these reporters, mainly because this is challenging and likely uninformative given that these are large and inherently will contain some heterogeneity. We will therefore in future studies focus on producing short sulfo-ST glycopeptides to perform structural studies using sulfated MAdCAM1 fragments in co-crystallization and/or cryo-EM experiments.

Our study also clearly indicates that multivalent interactions are involved in efficient binding of Siglec-7 and −15 to the identified mucin-like O-glycoproteins. Our binding assays with precomplexed Fc-chimera Siglecs showed increasing gradient-like reactivities with dST and CHST1-engineered sulfo-dST HEK293 cells expressing reporters with different mucin-like O-glycodomains (**Fig. 3**). In contrast, the reverse binding studies with isolated reporters and Siglec-expressing CHO cells revealed more clear-cut binding only with the sulfo-dST glycoforms of the MAdCAM1, CD43, and PSGL-1 reporters (**Fig. 5**). These results indicate a ligand affinity threshold for detection of binding on cells, which may be in agreement with previous work demonstrating a switch-like signaling response of Siglec-7, depending on ligand density and sialylation levels at the cell surface^29^. Moreover, a recent simulation study suggested receptor clustering on the membrane as a general mechanism for tuning the ligand binding threshold ^76^. Our findings support the notion that the natural dynamic state of Siglecs in the cell membrane may be important for multivalent binding to these select O-glycodomains and their spacing of O-glycan clusters, whereas the precomplexed Fc-chimera with four sialic acid binding sites may be more prone to bind high densities of the simple O-glycan epitope (**Fig. 1**). Of note, binding to such high-density epitopes overexpressed on cells might be further enhanced by fast rebinding rates due to the high local concentration of glycan epitopes (known as mass transport effect), as has been described for other lectin interactions ^77,78^. While there is evidence for clustering of Siglecs on the cell surface upon ligand binding ^29,50,51^, the exact oligomeric state of Siglec-7 is yet to be determined. However, to the best of our knowledge, the MAdCAM1, CD43, and PSGL-1 sulfo-ST reporters provide the first natural ligands demonstrated to interact with Siglecs on cells. These proteins are known to play important roles in immunity. MAdCAM1 is expressed on mucosal endothelial cells serving in recruitment of lymphocytes and a target for anti-inflammatory therapy ^79–82^, CD43 is expressed on most hematopoietic cells including T cells where its O-glycans characteristically change from core1 to core2 structures during T cell activation ^83^, and PSGL-1 is expressed on leukocytes serving as a key ligand for trafficking of immune cells and as an immune checkpoint for T cells ^84^. Whether Siglec interactions are involved in mediating these processes remains to be investigated.

Siglecs represent important immune checkpoints for therapeutic intervention ^12–14,85^, and Siglec-7 may be a promising target for enhancing therapeutic anti-tumor immunity with both inhibitor and antibody blocking strategies ^86^. Hence, studies of the cellular effects of engaging Siglec-7 with these complex reporters are needed. A challenge with probing large glycoproteins recombinantly expressed in mammalian cells for assaying Siglec signaling and bioactivities of immune cells is the down-stream processing with efficient removal of contaminants that non-specifically stimulate cells. In preliminary studies, we observed unspecific activation of monocytes by adding AF488-labeled mucins even in the absence of typical stimulants such as LPS (not shown), demonstrating the need for reliable workflows and assay formats that can define the functional effects of specific contextual glycoprotein binders. Knowledge of Siglec-7 signaling pathways, the induced cytokines, and downstream effects is still limited, and the few available reports mostly focused on global changes to sialoglycans displayed on a target cells ^50,87^. Here, we therefore did not explore the functional outcome on Siglec-7 signaling, but we demonstrated that sulfo-dST MAdCAM1 robustly induces Siglec-7 clustering and uptake. Sulfo-dST MAdCAM1, CD43, and PSGL-1 reporters may thus serve as natural ligand scaffolds to further dissect functional outcomes, e.g., by implementing recently developed Siglec signaling reporter cell lines ^29,88^. Such functional readouts will be essential for the rational design of selective inhibitors. Potent inhibitors of Siglecs based on sialic acid derivatization have been identified ^7,89–91^, and more recently, selective tetrameric Siglec-7/-9 inhibitors were generated targeting Siglec-7/9 to lysosomes ^92^. The scaffold for multivalent interactions embedded in human glycoprotein ligands such as MAdCAM1 may though provide for more natural and select inhibition.

In summary, our study provides strong evidence that Siglecs acquire selectivity and high-avidity in interactions with O-glycoproteins by multivalent binding to select clusters of sulfo-sialo O-glycans presented in repeated motifs found only on select O-glycoproteins. The results support the concept that clustered saccharide patches embedded in select proteins orchestrate Siglec interactions and their biological functions.

## MATERIALS & METHODS

Cell culture, detailed genetic engineering, mucin reporter sequences, mucin display cell-based arrays, mucin production and labeling, ELISA, sulfoglycomics, reverse CHO^Sig^ binding assays, internalization and immune cell binding assays, data analysis, and all additional information are available in the SI appendix.

### CRISPR/Cas9 targeted KO of GCNT1 and ST6GALNAC2-4

Glycosyltransferase gene KO in HEK cells was performed as described previously ^93^ using a library of validated gRNAs (glycoCRISPR) ^94^.

### Cell-binding assays of Siglec-7/15 to glycoengineered HEK cells

Human Siglec Fc chimera were precomplexed with goat anti-human IgG-Alexa647 at a 1:2 ratio for 15 min on ice. Cells were incubated with precomplexed Siglecs for 1 h on ice, washed, and analyzed by flow cytometry. For mucin display experiments, cells transiently expressed the respective FLAG-tagged transmembrane mucin reporters.

### Production and purification of mucin reporters

The secreted mucin reporters were stably expressed in glycoengineered HEK293 suspension cells and purified from the culture supernatant by DEAE-sepharose, eluting with increasing NaCl. Fractions containing the protein of interest were determined by SDS-PAGE.

### Binding of mucin reporters to CHO^Sig^

*CHO^Sig^* cells were incubated with AF488-labelled mucin reporters at the indicated concentrations for 1 h on ice. Siglec expression was co-stained with the respective anti-Siglec Ab for another 30 min on ice in the dark, before analysis by flow cytometry.

### Probing the human peripheral blood mononuclear cells (PBMCs) with flouro-labelled mucin reporters

Freshly isolated PBMCs from healthy donors were incubated with AF488-labelled mucin reporters for 1 h at 4°C, after Fc-receptor blocking for 10 min. Cells were co-stained with mouse-anti CD328/Siglec7, and optionally, mouse anti-human CD56, mouse anti-human CD14, and mouse anti-human CD3 for additional 30 min, before flow cytometric analysis.

## ACKNOWLEDGEMENTS

We thank James C. Paulson and Ajit Varki for valuable suggestions on the study and manuscript. Furthermore, we thank Lynn Janssen for technical assistance. This work was supported by the Lundbeck Foundation, the Novo Nordisk Foundation (NNF24OC0088218 to Y.N., NNF21OC0071658 to H.C., NNF22OC0073736 to R.L.M.), the Danish National Research Foundation (DNRF107), a Veni grant from the Dutch Research Council (NWO VI.Veni.202.045), and Academia Sinica (AS-IR-113-L04 to K.-H.K.). We would like to acknowledge Core Facility for Flow Cytometry and Single Cell Analysis and Core Facility for Integrated Microscopy, Faculty of Health and Medical Sciences, University of Copenhagen, and the Academia Sinica Common Mass Spectrometry Facilities for Proteomics and Protein Modification Analysis (AS-CFII-111-209) for part of the glycomics MS data acquisition. F.G. and T.J. were supported by the EMBO postdoctoral fellowship (ALTF 105-2023 and ALTF 336-2021, respectively).

## AUTHOR CONTRIBUTIONS

F.G., D.L.A.H.H., Y.C., D.d.W., T.J., M.J., V.T.C.V., S.F., H.T., K.K., R.L.M., and W.T.V.G. performed research and analyzed data. J.J., E.N.S., and M.S.M. provided CHO cell lines and recombinant Siglecs. F.G., G.M.J.B., H.C., C.B., and Y.N. designed experiments. F.G., H.C., and Y.N. conceived the project and wrote the manuscript with input from all authors.

## DATA AVAILABILITY

All study data are included in the article and supporting information.

