## Supplementary Information for "Siglec-7 and -15 recognize repeated clustered sulfo-sialo O-glycan motifs on select O-glycoproteins"

#### Content

- Supplementary Materials and Methods
- Supplementary Tables 1-3
- Supplementary Figures 1-11
- Supplementary References

#### Supporting Materials and Methods

*Cell culture*

HEK293 cells were cultured in DMEM + 10% FCS + 1x GlutaMAX and passaged every 2-3 days. $CHO^{Sig}$ cells were cultured in DMEM/F12 + 10% FCS + 1x GlutaMAX and passaged every 2-3 days.

*CRISPR/Cas9 targeted KO*

Briefly, one 6-well with cells at 70% confluency was transfected with 1 µg of gRNA in total (or multiple gRNAs for combinatorial KO) and 1 µg of plasmid encoding GFP-tagged Cas9 using lipofectamine 3000 (Invitrogen) following the manufacturer's protocol. One day later, GFP-positive cells were bulk-sorted and expanded for one week, before single-cell sorting into 96-well plates using a Sony SH800 cell sorter. Single-cell clones were expanded for two weeks and screened by Indel Detection by Amplicon Analysis (IDAA) (1). Final clones were validated by Sanger sequencing.

*Mucin TR reporters*

Constructs for transmembrane and secrete mucin tandem repeat and Glycocarrier reporters were previously described (2, 3). Codon-optimized dsDNA sequences of MAdCAM1-based reporter library were synthesized by IDT (gBlocks) and directly subcloned into transmembrane and secretion GFP-fusion vectors with BamHI and NotI sites (NEB). All tested reporter sequences were validated by Sanger sequencing and listed in **Tables S1-3**.

*Cell-binding assays of Siglec-7/15 to glycoengineered HEK cells*

Siglec Fc chimera (R&D) at the indicated concentrations were precomplexed with goat anti-human IgG-Alexa647 (Invitrogen) at a 1:2 ratio in PBA (PBS + 1% BSA) for 10-15 min on ice. Cells were detached using CellStripper (Corning) and $5x10^5$ cells were incubated in a 96-well plate with 50 µL of precomplexed Siglecs for 1 h on ice, washed three times with PBA, and analyzed on an LSR Fortessa III (BD). For mucin display experiments, cells transiently expressed the respective FLAG-tagged transmembrane mucin reporters (Table S1 for mucin sequences, construct design published previously (4)) by lipofectamine 3000 (Invitrogen) transfection one day before the experiment following the manufacturer's protocol, and mucin expression was quantified by GFP signal and anti-FLAG Ab-APC (BioLegend).

*Production and purification of mucin reporters*

The secreted mucin reporters were stably expressed in glycoengineered HEK293 cells selected by two weeks of culture in the presence of 1 µg/mL puromycin (Gibco). Cells were adapted for suspension culture by stepwise dilution into Freestyle F17 medium + 0.1% P188 + 2x GlutaMAX over the course of one week on an orbital shaker. A stable pool of cells was seeded at a density of 0.25-5 $x10^6$

cells/mL in 200 mL culture media for 5-7 days. Culture supernatant containing secreted mucin reporter was harvested by centrifugation and stored at -20°C until purification. Culture supernatant was filtered through a 0.45 μm filter and mixed 4:1 (v/v) with 5x buffer A (100 mM Tris-HCl, pH 7.5), and incubated overnight at 4°C with a DEAE-Sepharose resin slurry (Sigma) pre-equilibrated with buffer A. Resins were collected on gravity flow disposable plastic columns (Biorad) followed by washing with 5x column volumes (CV) of buffer A. Mucin reporter was eluted with 2CV of stepwise 100 mM NaCl increment (100 mM – 1 M NaCl) in buffer A. Fractions containing the protein of interest were determined by SDS-PAGE and desalted into water using Zeba spin desalting columns (ThermoFisher).

*ELISA*

ELISA assays were performed as described previously (4). Briefly, MaxiSorp 96-well plates (Nunc) were coated with serial dilutions of recombinant mucin reporters in carbonate-bicarbonate buffer (pH 9.6) overnight at 4°C, starting from 1 μg/mL. Plates were blocked with PLI-P buffer (0.5 M NaCl, 3 mM KCl, 1.5 mM $KH_2PO_4$, 6.5 mM $Na_2HPO_4$, 1% BSA, 1% Triton X-100, pH 7.4) for 1 h at RT. For Siglec binding experiments, plates were incubated for 1 h at RT with Siglec-7/15, precomplexed with anti-human Fc Ab-HRP (Merck) for 10 min. For PNA lectin experiments, mucin reporter coated plates were treated with neuraminidase at 20 mU/mL for 1 h at 37°C and incubated with biotinylated PNA (1 ug/mL, VectorLabs) for 1 h at RT followed by 1 h incubation with streptavidin-HRP (1:4000, Dako). In between all incubation steps, plates were washed with PBS-T. Plates were developed for 1-5 min with TMB substrate and quenched with 0.5 M $H_2SO_4$. Absorbance at 450 nm was measured using a BioTek plate reader (Agilent).

*Sulfoglycomics*

The O-glycans were released by beta-elimination and subsequently permethylated as described previously (5). Briefly, protein extracts from HEK cell lysates or purified mucin reporters were treated with 1 M sodium borohydride in 0.1 M sodium hydroxide at 45 °C. The released glycans were desalted and purified using a Dowex 50W-X8 resin and a C18 Sep-Pak cartridge (Waters). After the removal of borates by co-evaporation with 10% acetic acid in methanol under a nitrogen atmosphere, the reduced O-glycans were permethylated using a NaOH/DMSO slurry (50 mg NaOH powder, 200 μL dimethyl sulfoxide, and 200 μL iodomethane) at room temperature for 30 minutes with continuous vortexing. Permethylated glycans were then fractionated into pools of non-sulfated and sulfated glycans by eluting off an Oasis-MAX cartridge (Waters) with 95% acetonitrile, and 1 mM ammonium acetate in 80% acetonitrile, respectively. Then, permethylated O-glycans of HEK cell lysate were analyzed by LC-MS/MS on an Orbitrap Fusion as described (6). The permethylated non-sulfated glycans derived from the mucin reporters were dissolved in 50% methanol and spotted onto a target steel plate with 2,5-dihydroxybenzoic acid (DHB) matrix (Sigma-Aldrich) for analysis on an Autoflex

MALDI-TOF (Bruker) in positive ion mode with an m/z range of 300–1500. Data acquisition involved 5000 laser shots, and the glycan signals were manually annotated. Permethylated sulfated glycans were further desalted using a ZipTip (Merck Millipore) before injected directly into an Impact II Q-TOF (Bruker) for analysis in negative ion mode with an m/z range of 100–1500. The capillary voltage was set to 4500 V, with a dry nitrogen gas flow of 4 L/min at 180°C. The collision cell energy was set to 10 eV, and the collision RF was adjusted to 2500 Vpp. Data files were imported into Bruker Compass DataAnalysis 4.3 for manual annotation of sulfated glycans.

*Labeling of recombinant mucin reporters with Alexa488*

Lyophilized mucin reporters were dissolved into conjugation buffer (0.1 M $NaHCO_3$, pH 8.3) at 2.5 mg/mL and mixed in a 1:6 molar ratio with Alexa488-NHS ester (Lumiprobe) dissolved at 10 mg/mL in anhydrous DMSO, vortexed, and incubated for 1 h in the dark. Uncoupled dye was removed using Zeba dye and biotin removal spin columns (ThermoFisher), and the degree of labeling was monitored by measuring the absorbance at 280 and 519 nm using a nanodrop.

*Binding of mucin reporters to $CHO^{Sig}$*

*$CHO^{Sig}$* cells were detached with CellStripper (Corning) and 5 x$10^5$ cells were incubated in a 96-well plate with 50 µL of AF488-labelled mucin reporters at the indicated concentrations for 1 h on ice. Siglec expression was co-stained with the respective anti-Siglec Ab conjugated to APC or A700 (1:100, BioLegend) for another 30 min incubation on ice in the dark. Cells were washed thrice with PBA and analyzed using an LSR Fortessa III (BD).

*Internalization assay with MAdCAM1 reporter in $CHO^{Sig\text{-}7}$ cells*

$CHO^{Sig\text{-}7}$ were seeded onto 12 mm coverslips pre-coated for 20 min at RT with 500 µL PEI solution (25 µg/mL in 150 mM NaCl, sterile-filtered) at 25% confluency and grown overnight. After blocking with PBA for 1 h, cells were incubated with AF488-labelled mucin reporter for indicated time intervals at 37°C in the dark in a humidified chamber. Coverslips were washed with PBS and fixed with 4% paraformaldehyde for 10 min at RT. After three washes with PBS, coverslips were mounted using DAPI-containing Fluoromount G (ThermoFisher). and imaged using an LSM 900 (Zeiss) and a 63x/1.4 objective. All images were processed in Zen black (Zeiss).

*Probing the human peripheral blood mononuclear cells (PBMCs) with fluoro-labelled mucin reporters*

PBMCs were freshly isolated from blood samples of healthy donors (Sanquin) using density centrifugation over Ficoll-Paque (GE Healthcare). Cells were resuspended in 1x PBS containing 1% BSA (w/v) (PBA) and Fc-receptor blocking was performed using human TruStain FcX (Biolegend, #322301) for 10 min on ice. Subsequently, the PBMCs were washed with PBA and incubated with 10

μg/ml AF488-labelled mucin reporters diluted in PBA for 60 minutes at 4 °C. The cells were washed thrice with PBA and co-stained with AF700-conjugated mouse-anti CD328/Siglec7 (clone 6-434, Biolegend, #339209) for 30 minutes at 4 °C. Optionally, cells were stained with PE-conjugated mouseF. After three washes with PBA, the PBMCs were resuspended in PBA and analyzed using a CytoFLEX (Beckman Coulter) flow cytometer.
For experiments exploring the contribution of *cis*-inhibition, CD3-depleted PBMCs were treated with neuraminidase (from *C. perfringens*, J63259.MA, Sigma, 1:500) for 30 min on ice prior to incubation with 10 μg/ml AF488-labelled mucin reporters diluted in PBA for 60 min at 4 °C. To confirm removal of *cis*-ligands by neuraminidase, cells were stained with PNA-lectin (B1075, VectorLabs, 1:10,000) for 60 min on ice, washed twice with PBA and subsequentially stained with Alexa Fluor 647-conjugated streptavidin (S32357, Invitrogen, 1:2000) for 30 min on ice. Optionally, CD3-depleted PBMCs were co-stained with CD56-APC-Vio770 (REA196, Miltenyi, #130-114-548), CD14-Horizon-V450 (MoP9, BD, #560349), Siglec-7-APC (6-434, Biolegend, #339206), CD3-FITC (REA613, Miltenyi, #130-113-138) and Live/Dead® Fixable Aqua Dead Cell Stain Kit (Thermo Fisher Scientific, Cat. No. L34957) for 30 min on ice. Sample acquisition was performed on a BD FACS Canto II.

*Flow cytometry data analysis*

All FACS data was analyzed using FlowJo (BD Biosciences, version 10). Briefly, intact cells were selected in an SSC-A/FSC-A plot, followed by single-cell gating in a FSC-H/FSC-A plot. For mucin display experiments, mean fluorescent intensity (MFI) of the binding of Siglecs to GFP positive (expressing mucin TR reporters) and negative (not expressing) populations was quantified and compared. For $CHO^{Sig}$ binding assays, cells were gated for Siglec expression with anti-Siglec Ab. All multi-color staining results were compensated in FlowJo with single-stain controls. Detailed gating strategy for PBMC binding assays is outlined in Supplementary Fig. 11. MFI values were plotted using Prism 10 (GraphPad).

### Supplementary Table 1. Amino acid sequences of mucin reporters.

| Gene (HGNC) | Uniprot ID | # | Position in UniProt ID | length (aa) | Sequence |
|---|---|---|---|---|---|
| MUC1 | P15941 | | 121-260 | 140 | APDNKPAPGSTAPPAHGVTS APDTRPAPGSTAPPAHGVTS APDTRPAPGSTAPPAHGVTS APDTRPAPGSTAPPAHGVTS APDTRPAPGSTAPPAHGVTS APDTRPAPGSTAPPAHGVTS APDTRPAPGSTAPPAHGVTS |
| MUC2 | Q02817 | #1 | 1406-1554 | 152 | PSPPPTST TTLPPTTT PSPPTTTT TTPPPTTT PSPPITTT TTPPPTTT PSPPISTT TTPPPTTT PSPPTTT<br>PS PPTTT PSPPTTTT TTPPPTTT PS PPTTT PITPPAST TTLPPTTT PSPPTTTT TTPPPTTT PSPPTTT PITPPTST T |
| | | #2 | 1916-2064 | 152 | PTQTPTTTPITTTTTVTPTPTPT GTQTPTPTPITTTTTVTPTPTPT GTQTPTSTPITTTTTVTPTPTPT GTQTPTMTPITTTTTVTPTPTPT<br>GTQTPTTTPISTTTTVTPTPTPT GTQTPTSTPITTTTTVTPTPTPT GTQTPTTTPITTAA |
| MUC3A | Q02505 | #1 | 391-538 | 148 | TTEISSHSTPSFSSSTIYSTVSTSTTAISSLPPTSGTMVTSTTMTPSSLSTDIPFTTPTTITHHSVGSTGFLTTATDLTSTFTVSSSSAMSTSVIPSSPSIQNTETSSLVSMTSATTPNVRPTFVSTLSTPTSSLLTTFPATYSFSSS |
| | | #2 | 502-645 | 144 | MTSATTPNVRPTFVSTLSTPTSSLLTTFPATYSFSSSMSASSAGTTHTESISSPPASTSTLHTTAESTLAPTTTTSFTTSTTMEPPSTTAATTGTGQTTFTSSTATFPETTTPTPTTDMSTESLTTAMTSPPITSSVTSTNTVT |
| | | #3 | 612-762 | 151 | TTPTPTTDMSTESLTTAMTSPPITSSVTSTNTVTSMTTTTSPPTTTNSFTSLTSMPLSSTPVPSTEVVTSGTINTIPPSILVTTLPTPNASSMTTSETTYPNSPTGPGTNSTTEITYPTTMTETSSTATSLPPTSPLVSTAKTAKTPTTNL |
| MUC4 | E9PDY6 | | 1184-1327 | TQT | PLPVTSPSSASTGHAT PLLVTDTSSASTGHAT PLPVTDASSVSTDHAT SLPVTIPSAASTGHTT PLPVTDTSSASTGQAT<br>SLLVTDTSSVSTGDTT PLPVTSTSSASTGHVT PLHVTSPSSASTGHAT PLPVTSLSSASTGDTM |
| MUC5AC | P98088 | | 2708-2850 | 142 | TTSAPTT STTSAPTT STISAPTT STTSATTT STTSAPTP RRTSAPTT STISASTT STTSATTT STTSATTT<br>STISAPTT STTLSPTT STTSTTIT STTSAPIS STTSTPQT STTSAPTT STTSGPGT TSSPVPTT STTSAPTT |
| MUC6 | Q6W4X9 | #1 | 1786-1907 | 123 | TSATSSRLPTPFTTHSPPTGTTPISSTGPVTATSFQTTTTYPTPSHPHTTLPTHVPSFSTSLVTPSTHTVIIPTHTQMATSASIHSMPTGTIPPPTTIKATGSTHTAPPMTPTTSGTSQSPS |
| | | #2 | 1868-1953 | 86 | SIHSMPTGTIPPPTTIKATGSTHTAPPMTPTTSGTSQSPSSFSTAKTSTSLPYHTSSTHHPEVTPTSTTNITPKHTSTGTRTPVAH |
| MUC7 | Q8TAX7 | | 192-351 | 160 | PPTPSATTQAPPSSSAPPE TTAAPPTPPATTPAPPSSSAPPE TTAAPPTPSATTPAPLSSSAPPE TTAVPPTPSATTLDPSSASAPPE<br>TTAAPPTPSATTPAPPSSPAPQE TTAAPITTPNSSPTTLAPDTSET SAAPTHQTTTSVTTQTTTTKQPTSAP |
| OVGP1 (MUC9) | Q12889 | | 476-564 | 89 | AMTMTSVGHQSMTP GEKALTPVGHQSVTT GQKTLTSVGYQSVTP GEKTLTPVGHQSVTP VSHQSVSPGGTTMTP VHFQTETLRQNTVAP |
| MUC17 | Q685J3 | | 2181-2329 | 149 | LSTTPVDTSTPVTNSTEARSSPTTSEGTSMPTSTPSEGSTPFTSMPVSTMPVVTSEAST LSATPVDTSTPVTTSTEATSSPTTAEGTSIPTSTLSEGTTPLTSIPVSHTLVANSEVST<br>LSTTPVDSNTPFTTSTEASSPPPTAEGTSMP |
| MUC19 | Q7Z5P9 | | 3353-3501 | 149 | VTRTTRSSA GLTGKTGLSA GVTGKTGLSA EVTGTTRLSA GVTGTTGPSP GVTGTTGTPA GVTGTTELSA GVTGKTGLSS<br>EVTETTGLSY GVKRTIGLSA GSTGTSGQSA GVAGTTTLSA EVTGTTRPSA GVTGTTGLSA EVTEITGISA |
| MUC20 | Q8N307 | | 175-325 | 151 | ESSASSDSPHPVITPSRA SESSASSDGPHPVITPSRA SESSASSDGPHPVITPSRA SESSASSDGPHPVITPSRA<br>SESSASSDGPHPVITPSRA SESSASSDGPHPVITPSRA SESSASSDGPHPVITPSRA SESSASSDGPHPVITPSRA |
| MUC21 | Q5SSG8 | | 130-278 | 149 | SGASTATNSDSSTT SSGASTATNSDSSTT SSEASTATNSESSTT SSGASTATNSESSTV SSRASTATNSESSTT<br>SSGASTATNSESRTT SNGAGTATNSESSTT SSGASTATNSESSTP SSGAGTATNSESSTT SSGAGTATNSESSTV |
| MUC22 | E2RYF6 | | 336-483 | 148 | GTTTASMAG SETTVSTAG SETTTVSITG TETTMVSAMG SETTTNSTTS SETTVTSTAG SETTTVSTVG SETTTAYTAD<br>SETTAASTTG SEMTTVFTAG SETITPSTAG SETTTVSTAG SETTTVSTTG SETTTASTAH SETTAASTMG |
| MAdCAM1 | Q13477 | | 223-313 | 91 | VLHSPTSPEPPDTTSPESPDTTSPESPDTTSQEPPDTTSPEPPDKTSPEPAPQQGSTHTPRSPGSTRTRRPEISQAGPTQGEVIPTGSSKP |
| PSGL1 | Q14242 | #1 | 118-261 | 160 | AMEIQTTQPAATEAQTTQPVPTEAQTTPLAATEAQTTRLTATEAQTTPLAATEAQTTPPAATEAQTTQPTGLEAQTTAPAAMEAQTTAPAAMEAQTTPPAAMEAQTTQTTAMEAQTTAPEATEAQTTQPTATEAQTTPLAAMEA |
| | | #2 | 42-191 | 150 | QATEYEYLDYDFLPETEPPEMLRNSTDTTPLTGPGTPESTTVEPAARRSTGLDAGGAVTELTTELANMGNLSTDSAAMEIQTTQPAATEAQTTQPVPTEAQTTPLAATEAQTTRLTATEAQTTPLAATEAQTTPPAATEAQTTQPTGLEA |
| TIM1 | Q96D42 | #1 | 125-213 | 88 | PPKVTTTPIVTTVPTVTTVRTSTTVPTTTTVPMTTVPTTTVPTTMSIPTTTTVLTTMTVSTTTSVPTTTSIPTTTSVPVTTTVSTFVP |
|  |  | #2 | 125-213(Δ157-161) | 83 | PPKVTTTPIVTTVPTVTTVRTSTTVPTTTTVP-----TTTVPTTMSIPTTTTVLTTMTVSTTTSVPTTTSIPTTTSVPVTTTVSTFVP |
| TIM3 | Q8TDQ0 | | 131-201 | 71 | PAKVTPAPTRQRDFTAAFPRMLTTRGHGPAETQTLGSLPDINLTQISTLANELRDSRLANDLRDSGATIRI |
| TIM4 | Q96H15 | | 133-288 | 156 | RASTTTHRTATTTTRRTTTTSPTTTRQMTTTPAALPTTVVTTPDLTTGTPLQMTTIAVFTTANTCLSLTPSTLPEEATGLLTPEPSKEGPILTAESETVLPSDSWSSVESTSADTVLLTSKESKVWDLPSTSHVSMWKTSDSVSSPQPGASDTAVP |
| CD34 | P28906 | | 31-175 | 145 | MSLDNNGTATPELPTQGTFSNVSTNVSYQETTTPSTLGSTSLHPVSQHGNEATTNITETTVKFTSTSVITSVYGNTNSSVQSQTSVISTVFTTPANVSTPETTLKPSLSPGNVSDLSTTSTSLATSPTKPYTSSSPILSDIKAEI |
| CD43 | P16150 | | 20-243 | 224 | STTAVQTPTSGEPLVSTSEPLSSKMYTTSITSDPKADSTGDQTSALPPSTSINEGSPLWTSIGASTGSPLPEPTTYQEVSIKMSSVPQETPHATSHPAVPITANSLGSHTVTGGTITTNSPETSSRTSGAPVTTAASSLETSRGTSGPPLTMATVSLETSKGTSGPPVTMATDSLETSTGTTGPPVTMTTG<br>SLEPSSGASGPQVSSVKLSTMMSPTTSTNASTV |
| CD45 | P08575 | | 26-228 | 203 | QSPTPSPTGLTTAKMPSVPLSSDPLPTHTTAFSPASTFERENDFSETTTSLSPDNTSTQVSPDSLDNASAFNTTGVSSVQTPHLPTHADSQTPSAGTDTQTFSGSAANAKLNPTPGSNAISDVPGERSTASTFPTDPVSPLTTTLSLAHHSSAALPARTSNTTITANTSDAYLNASETTTLSPSGSAVIST<br>TTIATTPSKPTC |
| CD68 | P34810 | | 29-154 | 126 | SATLLPSFTVTPTVTESTGTTSHRTTKSHKTTTHRTTTTGTTSHGPTTATHNPTTTSHGNVTVHPTSNSTATSQGPSTATHSPATTSHGNATVHPTSNSTATSPGFTSSAHPEPPPPSPSPSPTSK |
| SDC1 | P18827 | | 56-220 | 165 | TLSQQTPSTWKDTQLLTAIPTSPEPTGLEATAASTSTLPAGEGPKEGEAVVLPEVEPGLTAREQEATPRPRETTQLPTTHLASTTTATTAQEPATSHPHRDMQPGHHETSTPAGPSQADLHTPHTEDGGPSATERAAEDGASSQLPAAEGSGEQDFTFETSGENT |
| SDC3 | O75056 | | 139-303 | 165 | ATSPLVVTEVPEEPSQRATTVSTTMATTAATSTGDPTVATVPATVATATPSTPAAPPFTATTAVIRTTGVRRLLPLPLTTVATARATTPEAPSPPTTAAVLDTEAPTPRLVSTATSRPRALPRPATTQEPDIPERSTLPLGTTAPGPTEVAQTPTPETFLTTIRD |

**Supplementary Table 2. Amino acid sequences of Glycocarrier reporters.**

| Glycocarrier | Length (aa) | Sequence |
|---|---|---|
| 1xThr | 198 | AEAAA**T**PAPAKAEAAA**T**PAPAKAEAAA**T**PAPAKAEAAA**T**PAPAKAEAAA**T**PAPAKAEAAA**T**PAPAKAEAAA**T**PAPAKAEAAA**T**PAPAKAEAAA**T**PAPAKAEAAA**T**PAPAKAEAAA**T**PAPAKAEAAA**T**PAPAKAEAAA**T**PAPAKAEAAA**T**PAPAKAEAAA**T**PAPAKAEAAA**T**PAPAKAEAAA**T**PAPAKAEAAA**T**PAPAK |
| 2xThr | 216 | AEAAA**TT**PAPAKAEAAA**TT**PAPAKAEAAA**TT**PAPAKAEAAA**TT**PAPAKAEAAA**TT**PAPAKAEAAA**TT**PAPAKAEAAA**TT**PAPAKAEAAA**TT**PAPAKAEAAA**TT**PAPAKAEAAA**TT**PAPAKAEAAA**TT**PAPAKAEAAA**TT**PAPAKAEAAA**TT**PAPAKAEAAA**TT**PAPAKAEAAA**TT**PAPAKAEAAA**TT**PAPAKAEAAA**TT**PAPAKAEAAA**TT**PAPAK |
| 4xThr | 224 | AEAAA**TTTT**PAPAKAEAAA**TTTT**PAPAKAEAAA**TTTT**PAPAKAEAAA**TTTT**PAPAKAEAAA**TTTT**PAPAKAEAAA**TTTT**PAPAKAEAAA**TTTT**PAPAKAEAAA**TTTT**PAPAKAEAAA**TTTT**PAPAKAEAAA**TTTT**PAPAKAEAAA**TTTT**PAPAKAEAAA**TTTT**PAPAKAEAAA**TTTT**PAPAKAEAAA**TTTT**PAPAKAEAAA**TTTT**PAPAKAEAAA**TTTT**PAPAK |
| 6xThr | 192 | AEAAA**TTTTTT**PAPAKAEAAA**TTTTTT**PAPAKAEAAA**TTTTTT**PAPAKAEAAA**TTTTTT**PAPAKAEAAA**TTTTTT**PAPAKAEAAA**TTTTTT**PAPAKAEAAA**TTTTTT**PAPAKAEAAA**TTTTTT**PAPAKAEAAA**TTTTTT**PAPAKAEAAA**TTTTTT**PAPAKAEAAA**TTTTTT**PAPAKAEAAA**TTTTTT**PAPAK |
| cluster1 | 209 | AEAAGAPAPAKFAEAA**T**PAPAKFAEAA**S**PAPAKFAEAA**T**PAPAKFAEAA**S**PAPAKFAEAA**T**PAPAKFAEAA**S**PAPAKFAEAA**T**PAPAKFAEAA**S**PAPAKFAEAA**S**PAPAKFAEAA**T**PAPAKFAEAA**S**PAPAKFAEAA**T**PAPAKFAEAA**S**PAPAKFAEAA**T**PAPAKFAEAA**S**PAPAKFAEAA**T**PAPAKFAEAA**S**PAPAKFAEAA**T**PAPAK |
| cluster2 | 82 | AEAAVPAPAKFAEAA**T**PAPAKFAAEAA**ST**PAPAKFAEAA**TT**PAPAKFAEAA**SS**PAPAKFGAEAA**TS**PAPAKFAEAA**T**PAPAK |
| cluster3 | 145 | AEAAGPAPAKFAEAA**T**PAPAKFLAEAA**STT**PAPAKFAAEAA**STS**PAPAKFPAEAA**TTT**PAPAKFAEAA**SSS**PAPAKFALAEAA**TST**PAPAKFGAEAA**TSS**PAPAKFAPAEAA**TTS**PAPAKFVAEAA**SST**PAPAKFAEAA**T**PAPAK |
| cluster4#1 | 170 | AEAAAPAPAKFAEAA**T**PAPAKFPAEAA**TTTT**PAPAKFGAEAA**TSSS**PAP AEAA**TSTTTT**PAPA AKFVAAEAA**TTST**PAPAKFLAEAA**TSST**PAPAKFGAAEAA**STTT**PAPAKFVAEAAA**SSTS**PAPAKFVLAEAA**STST**PAPAKFGAEAA**TTSS**PAPAKFAEAA**T**PAPAK |
| cluster4#2 | 157 | AEAAPPAPAKFAEAA**T**PAPAKFAEAA**SSSS**PAPAKFAPAEAA**TTTS**PAPAKFLAEAA**TSTT**PAPAKFVAAEAA**SSST**PAPAKFVAEAA**TSTS**PAPAKFGAAEAAA**STTS**PAPAKFGLAEAA**SSTT**PAPAKFGAAEAA**STSS**PAPAKFAEAA**T**PAPAK |
| cluster5 | 118 | AEAAPPAPAKFAEAA**T**PAPAKFLEAAA**TTTTT**SPAPAKFKFGAEAA**TTSTTT**PAPAKFLAEAA**TTTSTT**PAPAKFVEAA**TTTTST**PAPAKFPAEAA**STTTTT**PAPAKFAEAA**T**PAPAK |

**Supplementary Table 3. Amino acid sequences of MAdCAM1-based reporters.**

| Name | Length (aa) | Sequence |
|---|---|---|
| Full-length | 91 | VLHSPTSPEPPDTTSPESPDTTSPESPDTTSQEPPDTTSPEPPDKTSPEPAPQQGSTHTPRSPGSTRTRRPEISQAGPTQGEVIPTGSSKP |
| Fragment1 | 52 | VLHSPTSPEPPDTTSPESPDTTSPESPDTTSQEPPDTTSPEPPDKTSPEPAP |
| Fragment2 | 41 | QQGSTHTPRSPGSTRTRRPEISQAGPTQGEVIPTGSSKP |
| 4/6/7A | 52 | VLHAPAAPEPPDTTSPESPDTTSPESPDTTSQEPPDTTSPEPPDKTSPEPAP |
| 13-15A | 52 | VLHSPTSPEPPDAAAPESPDTTSPESPDTTSQEPPDTTSPEPPDKTSPEPAP |
| 21-23A | 52 | VLHSPTSPEPPDTTSPESPDAAAPESPDTTSQEPPDTTSPEPPDKTSPEPAP |
| 29-31A | 52 | VLHSPTSPEPPDTTSPESPDTTSPESPDAAAQEPPDTTSPEPPDKTSPEPAP |
| 37-39A | 52 | VLHSPTSPEPPDTTSPESPDTTSPESPDTTSQEPPDAAAPEPPDKTSPEPAP |
| Space13 | 52 | VLHSPTSPEPPDAAAPESPDTTSPESPDAAAQEPPDTTSPEPPDAAAPEPAP |
| Space21 | 52 | VLHSPTSPEPPDAAAPESPDAAAPESPDTTSQEPPDAAAPEPPDAAAPEPAP |
| Space29 | 52 | VLHSPTSPEPPDAAAPESPDAAAPESPDAAAQEPPDTTSPEPPDAAAPEPAP |
| Space37 | 52 | VLHSPTSPEPPDAAAPESPDAAAPESPDAAAQEPPDAAAPEPPDKTSPEPAP |
| 2xclusters | 52 | VLHAPAAAPEPPDAAAPEAPDAAAPEAPDAAAQEPPDTTSPEPPDKTSPEPA |
| 3xclusters | 52 | VLHSPTSPEPPDTTSPESPDTTSPESPDAAAQEPPDAAAPEPPDAAAPEPAP |
| Uncharged | 52 | VLHSPTSPAPPATTSPASPATTSPASPATTSQAPPATTSPAPPAATSPAPAP |
| repeatPx6 | 48 | PDTTSPEPPDTTSPEPPDTTSPEPPDTTSPEPPDTTSPEPPDTTSPEP |
| repeatSx6 | 48 | PDTTSPESPDTTSPESPDTTSPESPDTTSPESPDTTSPESPDTTSPEP |

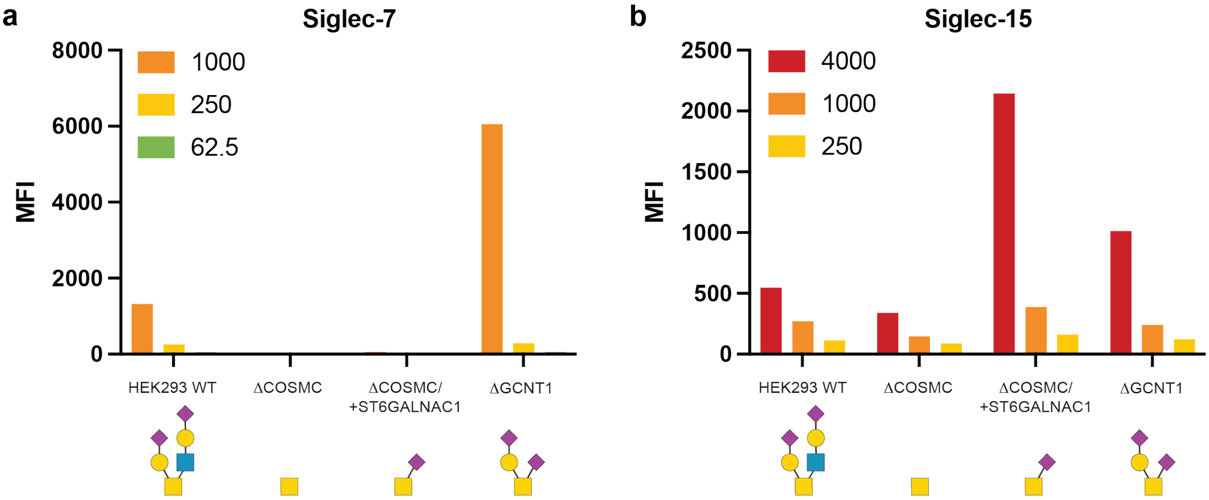


**Supplementary Fig. 1. Genetic dissection of Siglec-7/15 binding to non-sulfated O-glycans on HEK293 cells**. Bar diagrams show flow cytometry analysis (mean fluorescence intensity, MFI) of Siglec-7 (**a**) or Siglec-15 (**b**) binding to glycoengineered HEK293 cells with KO/KI of glycosyltransferases as indicated. Siglec-7/15 were precomplexed with anti-human-IgG antibody-Alexa647 (Invitrogen A-21445) and titrated to cells at the indicated concentrations in ng/mL. Data presented here is the same data as in Figure 2 but excluding the CHST1 KI cell lines to increase the resolution in the low MFI range.

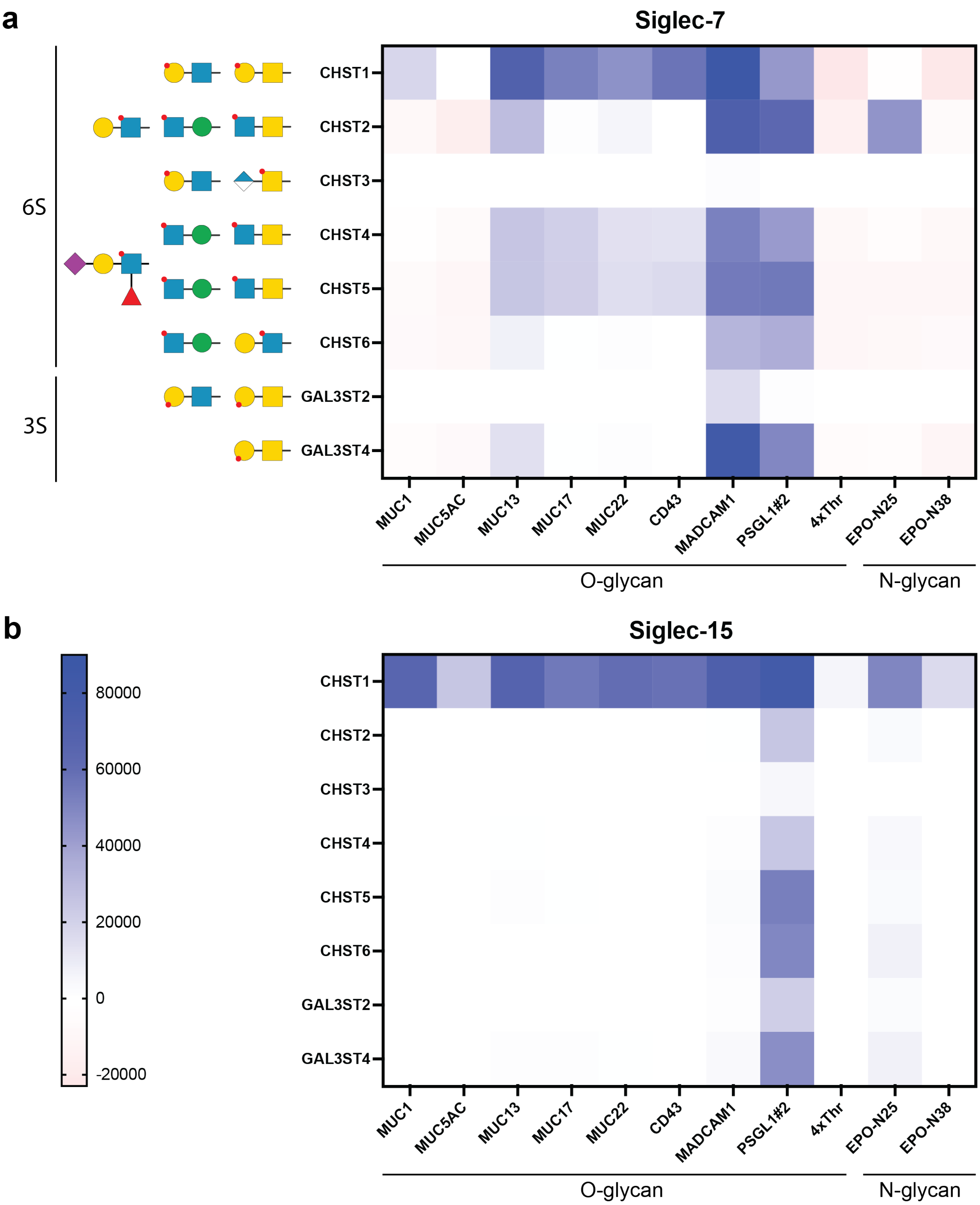


**Supplementary Fig. 2. Flow cytometry analysis of Siglec-7/15 binding to a sulfotransferase KI library of HEK293 cells expressing mucin reporters.** Mucins were expressed on the surface of HEK293 cells with different sulfotransferase KIs enabling 6-O- or 3-O-sulfation. (**a**) Heat map displaying binding increase (ΔMFI) of Siglec-7 upon mucin expression. (**b**) Heat map displaying binding increase of Siglec-15 upon mucin expression. Siglec-7/15 (at 1 µg/mL) were pre-complexed with anti-human-IgG antibody-Alexa647 (Invitrogen A-21445) prior to incubation with cells. Representative MFI values from two independent experiments.

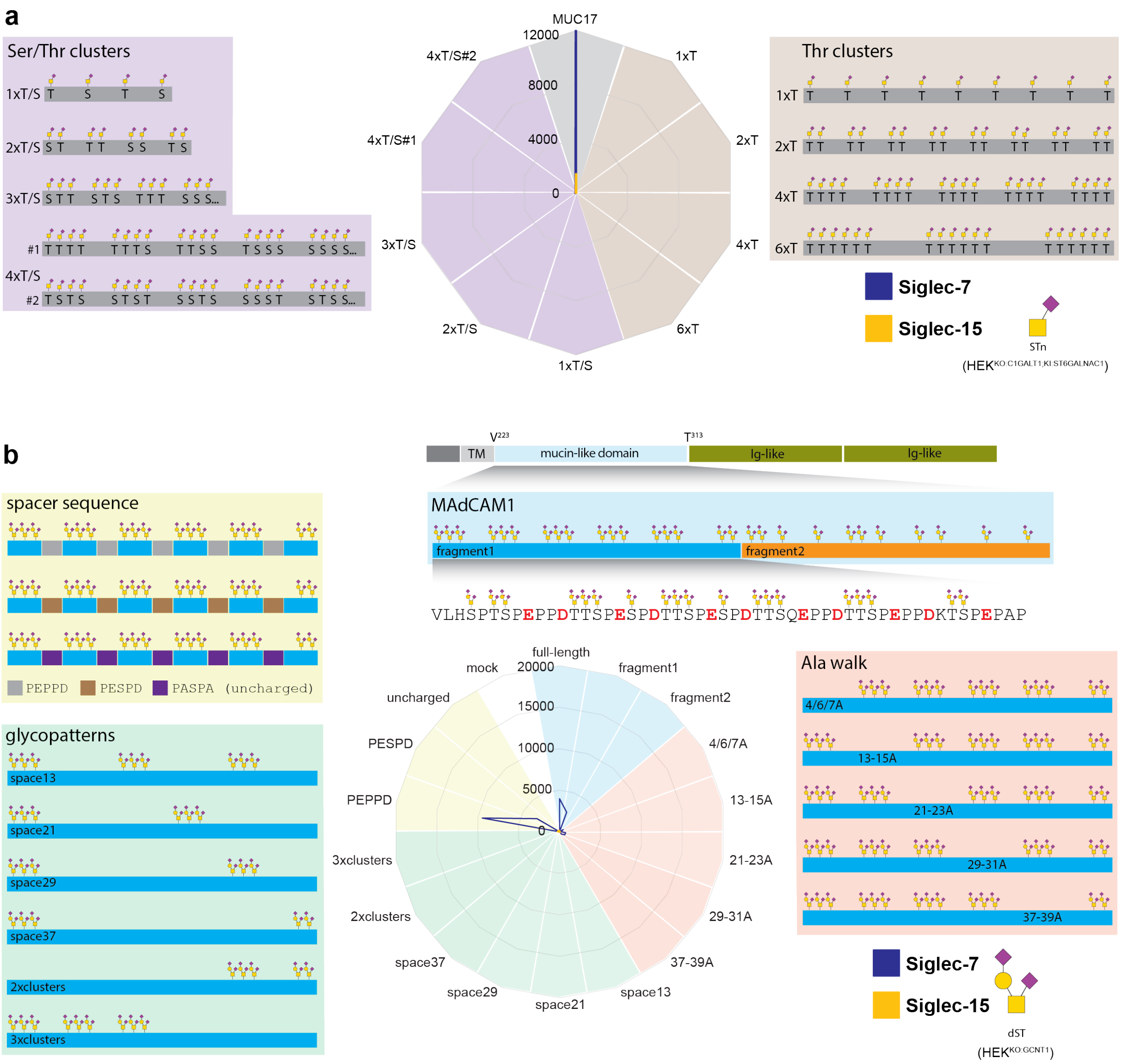


**Supplementary Fig. 3. Flow cytometry analysis of putative O-glycan cluster motifs for Siglec-7/15 binding by Glycocarriers and MAdCAM1-based reporters.** (**a**) Binding of Siglec-7 (blue) and Siglec-15 (yellow) (at 1 µg/mL, pre-complexed with anti-human-IgG antibody-Alexa647) to a library of rationally designed O-Glycocarriers (2) displayed on cells engineered to produce STn O-glycans (HEK$^{KO:C1GALT1;KI:ST6GALNAC1}$) as indicated. Color code of radar plots: Brown, clusters of 1-5 Thr repeats; Purple, clusters of 1-4 Ser/Thr repeats. Representative MFI values from two independent experiments. (**b**) Binding of Siglec-7 and Siglec-15 to a library of MAdCAM1-based reporters displayed on dST cells (HEK$^{KO:GCNT1}$). Color code of radar plots: Blue, MAdCAM1 fragments; Red, alanine walk mutants (removal of individual TTS repeats); Green, O-glycan cluster mutants (deletion of TTS repeats); Yellow, Spacer mutants. Cell surface expression of the Glycocarriers and MAdCAM1 reporters was verified with an anti-FLAG mAb. Mock-transfected cells were subjected to the same transfection protocol but leaving out DNA. Representative MFI values from four independent experiments.

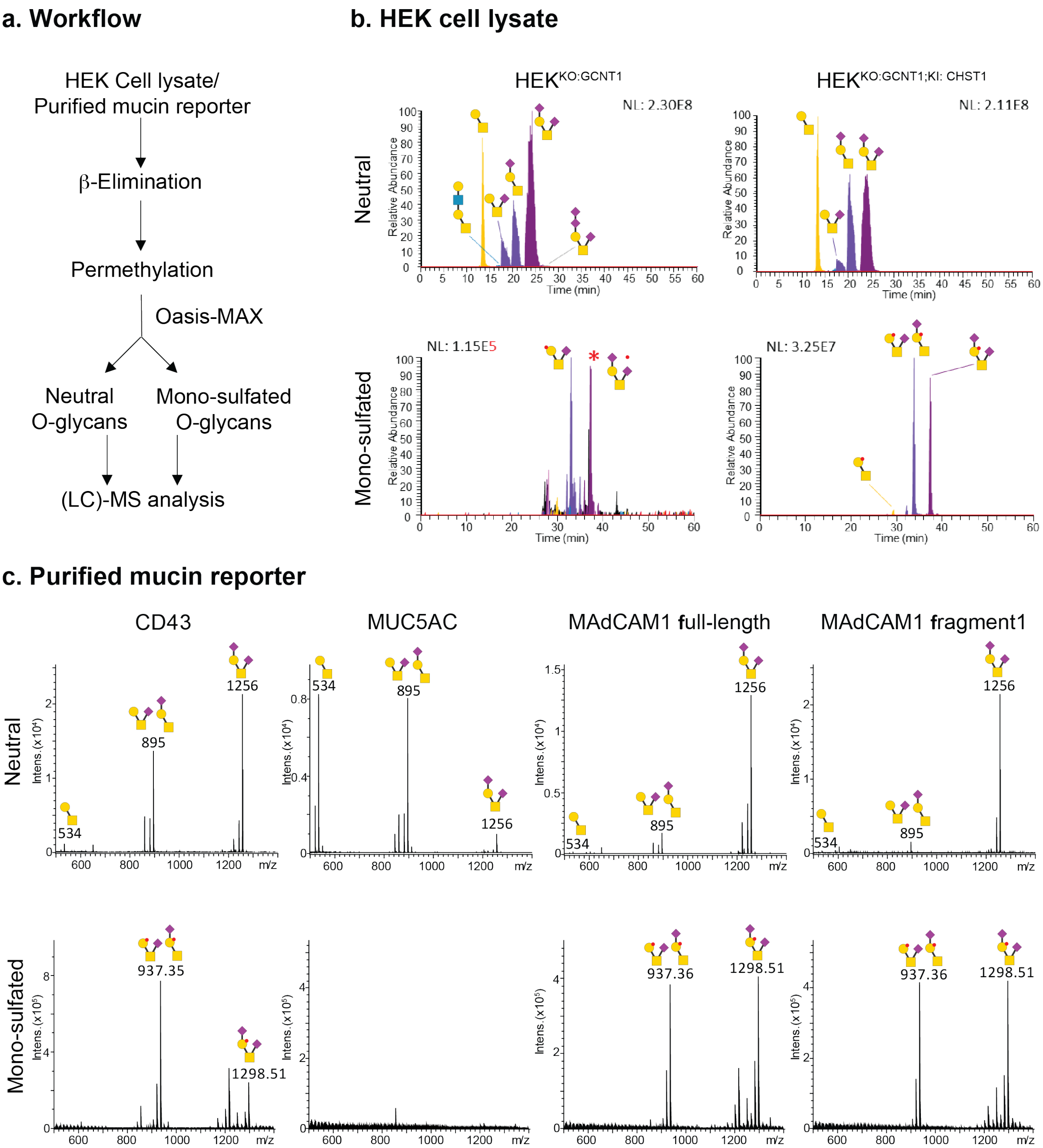


**Supplementary Fig. 4. Identification of O-glycans derived from HEK$^{KO:GCNT1;KI:CHST1}$ cell lysate and purified mucin reporters by mass spectrometry.** (**a**) Workflow for analyzing the O-glycans based on (5). O-glycans were released through β-elimination, permethylated, separated using an Oasis-MAX cartridge and analyzed by (LC)-MS. (**b**) Extracted ion chromatograms from LC-MS/MS analysis of the neutral O-glycans (upper panel) and sulfated O-glycans (lower panel) from HEK cell lysates. (**c**) MS profiles of non-sulfated glycans from the reporters in positive-ion mode (upper panel) and sulfated glycans in negative-ion mode (lower panel). Secreted reporters CD43, MUC5AC, MAdCAM1 full-length, and MAdCAM1 fragment1 were expressed in HEK$^{KO:GCNT1;KI:CHST1}$ and purified.

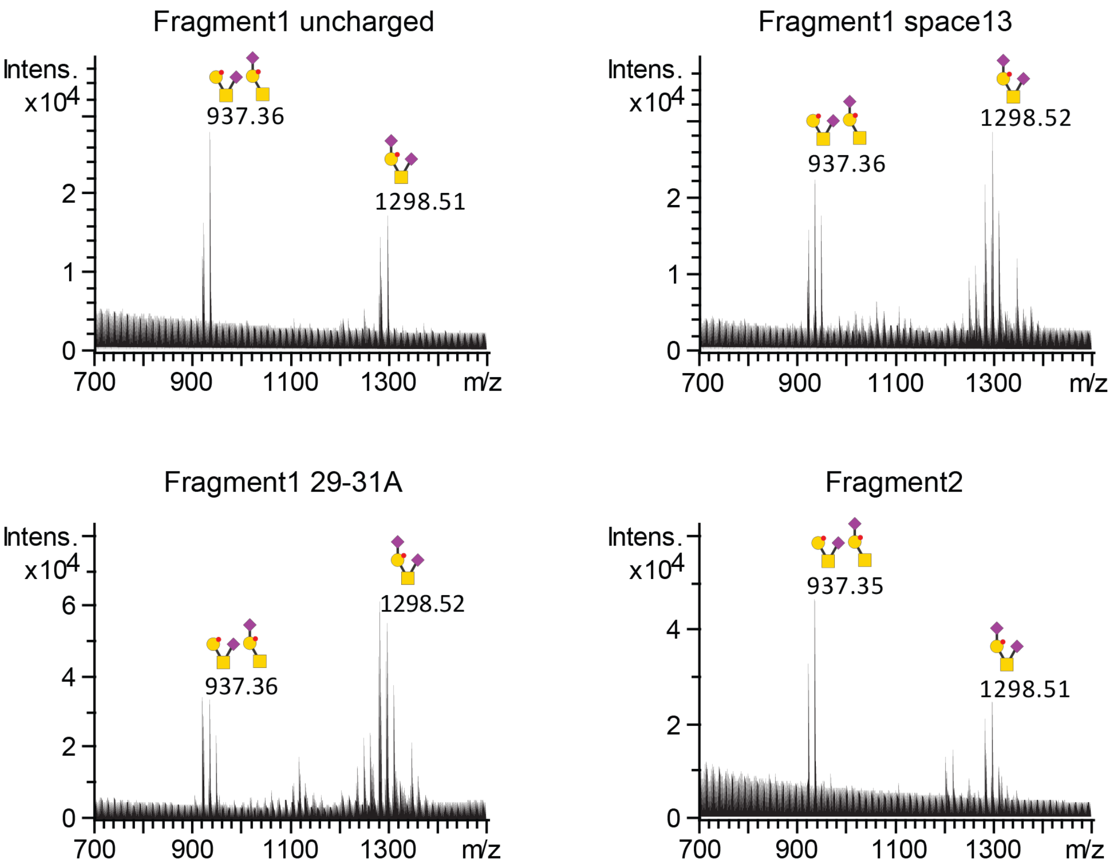


**Supplementary Fig. 5. Identification of sulfated O-glycans derived from purified MAdCAM1-based reporters by mass spectrometry.** MS profiles of sulfated glycans in negative-ion mode. Secreted reporters were expressed in HEK$^{KO:GCNT1;KI:CHST1}$ and purified, and the presence of non-sulfated sialoglycans (mST/dST) was confirmed in positive-ion mode. Note that all tested MAdCAM1-derived reporters display sulfo-mST and sulfo-dST structures.

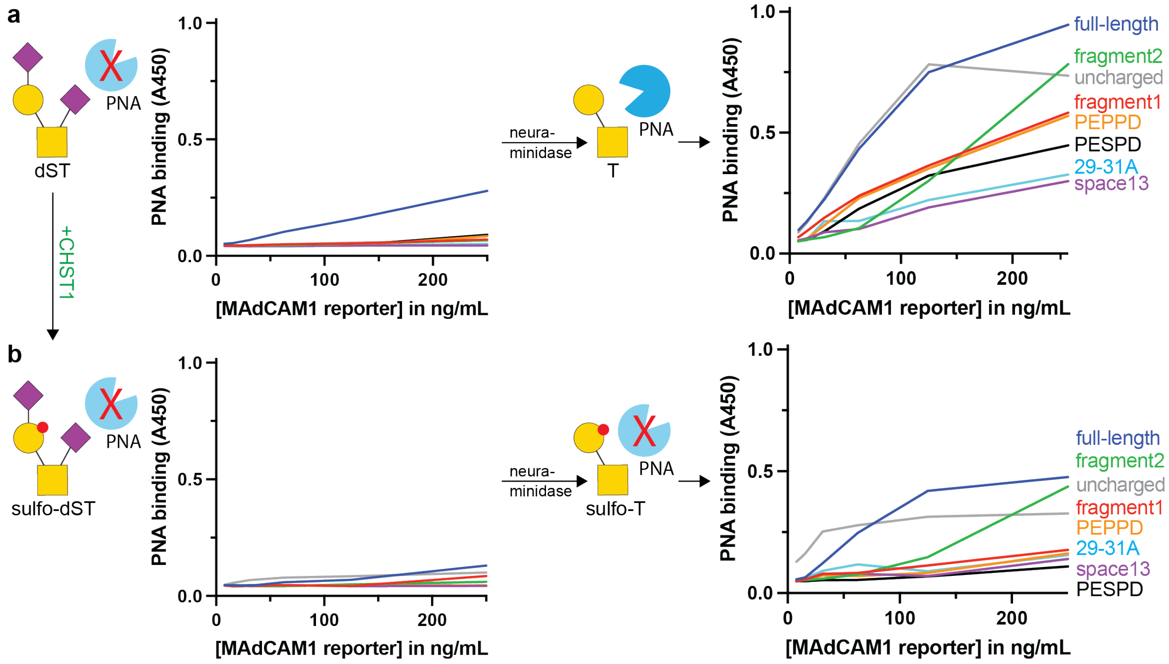


**Supplementary Fig. 6. ELISA analysis of PNA binding to secreted, purified, neuraminidase-treated MAdCAM1 reporters.** Plots display PNA binding at different concentrations of MAdCAM1-based reporters purified from (**a**) HEK$^{KO:GCNT1}$ (capacity for dST) or (**b**) HEK$^{KO:GCNT1;KI:CHST1}$ (capacity for sulfo-dST). Plots represent mean values from N ≥ 3.

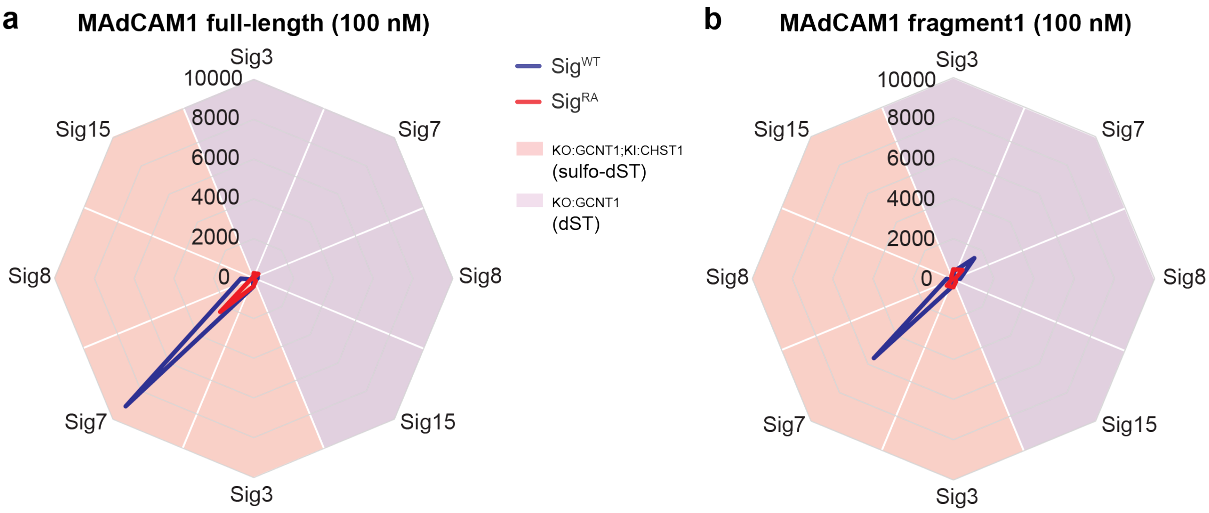


**Supplementary Fig. 7. Flow cytometry analysis of binding to CHO cells displaying different Siglecs.** (**a**) Binding of non-sulfated (purple) or sulfated (salmon) MAdCAM1 to CHO cells displaying different Siglecs, either as wildtype (blue) or as inactive RA mutant (red). MFI values represent mean from three independent experiments. (**b**) Binding of non-sulfated (purple) or sulfated (light red) MAdCAM1 fragment1 to CHO cells displaying different Siglecs, either as wildtype (blue) or as inactive RA mutant (red). Representative MFI values from two independent experiments.

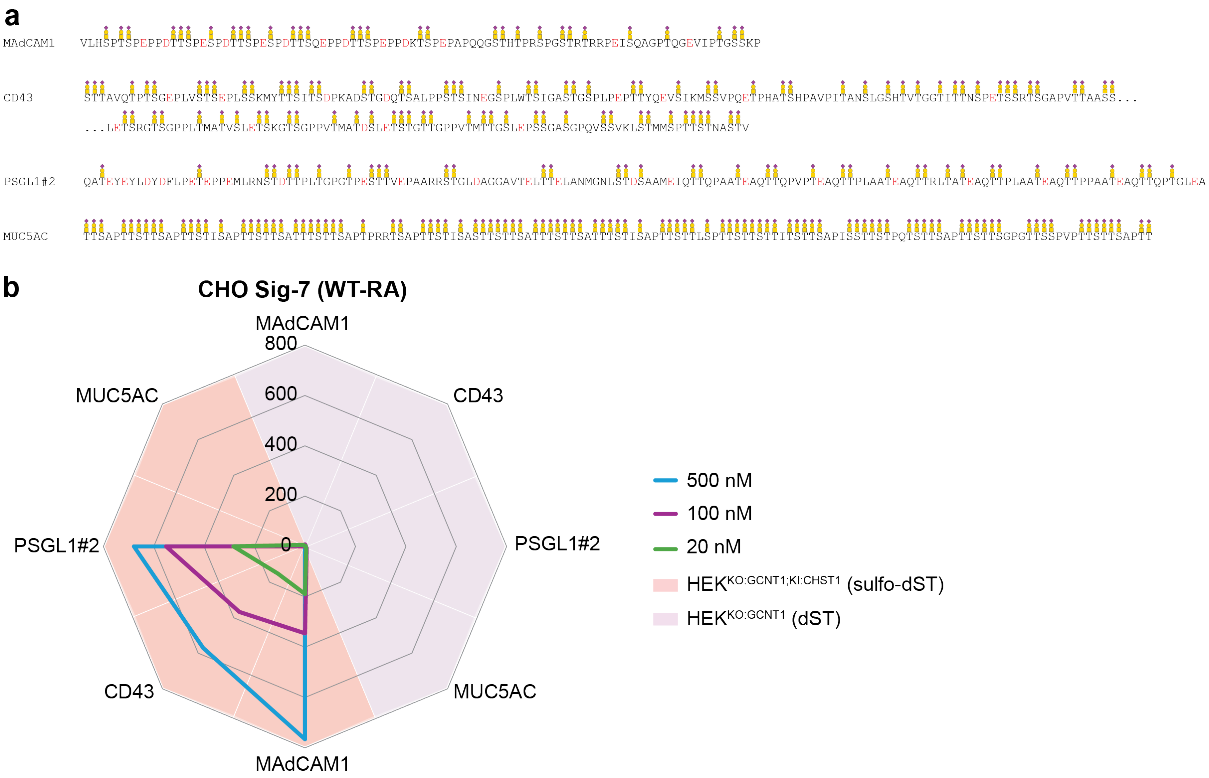


**Supplementary Fig. 8. Flow cytometry analysis of binding of different mucins to Siglec-7 installed on CHO cells.** (**a**) Mucin reporters probed with CHO$^{Sig\text{-}7}$. O-glycosylation sites and acidic residues (red) are marked. (**b**) Binding of non-sulfated (purple) vs. sulfated (salmon) mucins to CHO$^{Sig\text{-}7}$ cells at indicated concentrations. Values represent MFI obtained by subtracting MFI(CHO$^{Sig\text{-}7R124A}$) from MFI (CHO$^{Sig\text{-}7WT}$), reproduced in three independent experiments.

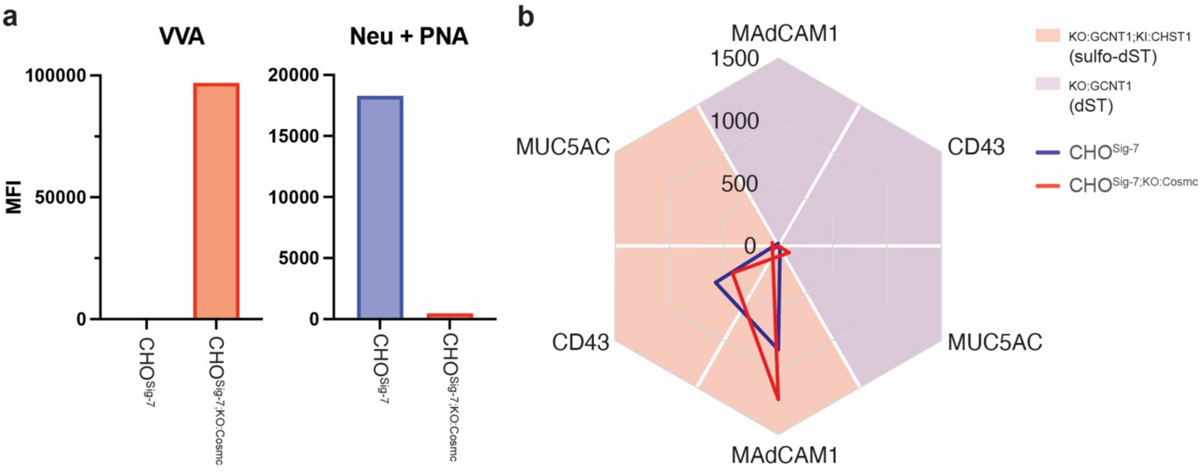


**Supplementary Fig. 9. Flow cytometry analysis of binding of different mucins to Siglec-7 installed on CHO$^{\text{Sig-7;KO:COSMC}}$.** (**a**) Validation of the COSMC KO cell line with VVA lectin (binds Tn, left panel) and PNA lectin (binds core1, right panel). For PNA staining, cells were pre-treated with neuraminidase (Neu) to expose potential (sialylated) core1 structures. Note that CHO$^{\text{Sig-7;KO:COSMC}}$ VVA but not PNA staining suggesting complete truncation of O-glycans (Tn). (**b**) Binding of non-sulfated (purple) vs. sulfated (salmon) mucins to CHO$^{\text{Sig-7}}$ (blue) or CHO$^{\text{Sig-7;KO:COSMC}}$ (red). Note that binding profiles are very similar suggesting only limited/no masking by *cis*-ligands.

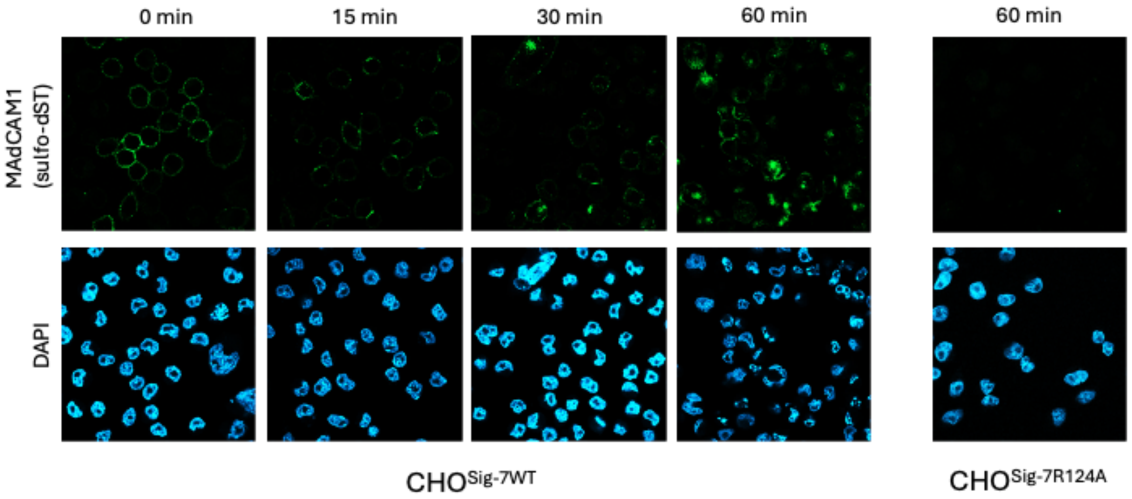


**Supplementary Fig. 10. Confocal microscopy imaging of MAdCAM1 binding to and uptake by CHO cells displaying Siglec-7.** Representative images of CHO$^{\text{Sig-7WT}}$ cell stainings with sulfated MAdCAM1-Alexa488 (green) and DAPI (blue) after incubation at 37°C for indicated time intervals. Right panel shows staining of CHO$^{\text{Sig-7R124A}}$ after 60 min for comparison.

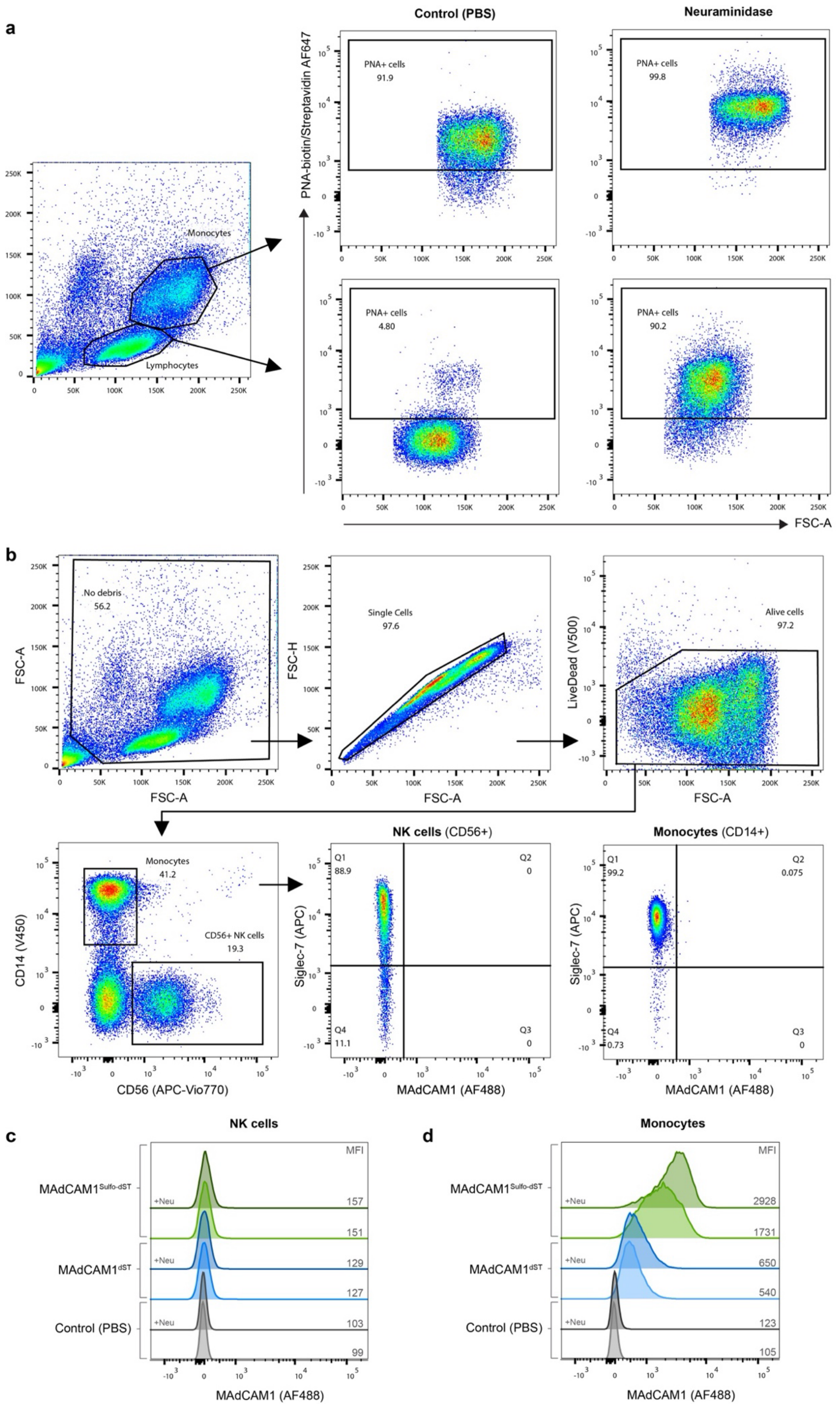


**Supplementary Fig. 11. Flow cytometry analysis of sulfo-dST MAdCAM1 binding to primary human NK cells and monocytes expressing Siglec-7.** (legend continues next page)

(…Supplementary Fig. 11 continued) (**a**) Neuraminidase treatment of PBMCs. PNA lectin (y axis) is used to detect exposed core1 O-glycans after sialic acid removal on monocyte and lymphocyte populations. (**b**) Gating strategy for isolating NK cell and monocyte populations. (**c**) AF488-labeled MAdCAM1 (dST vs. sulfo-dST) binding to NK cells with or without neuraminidase pre-treatment. (**d**) AF488-labeled MAdCAM1 (dST vs. sulfo-dST) binding to monocytes with or without neuraminidase pre-treatment.
